# Diabetes and Immunosuppression Drive Distinct Patterns of *Candidozyma auris* Skin Colonization and Dissemination in Mice

**DOI:** 10.64898/2026.05.24.727467

**Authors:** Kaustav Das Gupta, Derek Quintanilla, Eman G. Youssef, Kavya Gupta, Ashraf S. Ibrahim, Shakti Singh

## Abstract

*Candidozyma auris* is an emerging, multidrug-resistant (MDR) fungal pathogen that persistently colonizes human skin and disproportionately causes invasive infections in patients with metabolic or immune dysfunction. Despite strong epidemiological links to diabetes and immunosuppression, how these host conditions shape skin colonization, immune defense, and dissemination remain poorly defined. Here, we establish complementary murine model of C. auris skin colonization under immunocompetent, immunosuppressed, and diabetic ketoacidosis (DKA) conditions. DKA mice exhibited significantly increased skin fungal burden, impaired clearance, and frequent systemic dissemination. Notably, DKA permitted dissemination despite preserved granulocyte and neutrophil recruitment to the skin, indicating a functional rather than quantitative defect in innate immunity. Consistent with this, phagocytes from DKA mice displayed impaired antifungal activity characterized by reduced phagocytosis and killing despite elevated reactive oxygen species production. Hyperglycemic and ketone-rich conditions (BHB) remodel the *C. auris* cell wall, reducing mannan, increasing chitin, and upregulating adhesins, thereby enhancing adhesion and inflammatory activation while impairing neutrophil killing. Together, these findings reveal host metabolic dysfunction as a primary driver of persistent *C. auris* skin colonization and dissemination, identify qualitative defects in innate antifungal immunity as a key determinant of invasive risk and highlight metabolic condition as a critical target for infection prevention strategies.

**Short Summary:** This study establishes the first physiologically relevant murine model of *Candidozyma auris* skin colonization under diabetic ketoacidosis and immunosuppression, revealing distinct immune dysfunction and systemic dissemination that can inform targeted antifungal strategies.

## INTRODUCTION

*Candidozyma* auris (synonym *Candida auris*) is an emerging multidrug-resistant (MDR) fungal pathogen with a unique capacity for persistent skin colonization and nosocomial transmission, reported in more than 55 countries worldwide^1–3^. Its extensive antifungal resistance and high mortality (up to ∼60%) in invasive infections have led the World Health Organization (WHO) and Centers for Diseases Control and Prevention (CDC) to classify *C. auris* as a critical priority pathogen and an urgent threat, respectively^4–9^. A defining feature of *C. auris* is its exceptional capacity to adhere to and persist on both biotic and abiotic surfaces, including human skin and healthcare fomites^10,11^. Experimental studies have demonstrated robust biofilms under axillary-like skin conditions and on porcine skin, conferring tolerance to desiccation and resistance to commonly used disinfectants^12–16^. These properties enable prolonged colonization and facilitate transmission within healthcare environments.

Clinically, *C. auris* skin colonization is disproportionately observed in patient with immunosuppression or poorly controlled diabetes and is strongly associated with an increased risk of subsequent disseminated diseases and candidemia^17–25^. Asymptomatic colonization is often prolonged, enabling colonized individuals to contaminate their surroundings and drive nosocomial spread^3,26–32^. Despite these strong epidemiological associations, it remains unclear whether immune suppression and metabolic dysfunction promote *C. auris* persistence through similar or fundamentally distinct mechanisms. Whether diabetes primarily increases susceptibility by limiting immune cell recruitment or by impairing antifungal effector function has not been resolved.

Recent studies have identified fungal adhesins such as Agglutinin-like sequence (ALS) protein family^33,34^, Hyphal-regulated/Iff-like (HIL) protein family^35,36^ and novel surface colonization factor-1 (SCF-1)^37^. These adhesins and their gene amplification, particularly ALS4^38^ and biofilm-associated factors, have been implicated in *C. auris* adherence to abiotic and biotic surfaces ^33–35,37,39^. Together these fungal factors and altered local immune signatures are contributors to *C. auris* skin persistence, with immune responses distinct from those elicited by *Candida albicans*. Murine and porcine skin models have shown altered neutrophil, monocyte, and lymphoid responses and have demonstrated the potential for dissemination from cutaneous sites^40–42^. However, most existing *in vivo* models focus on immunocompetent or immunosuppressed hosts and do not directly interrogate the impact of metabolic dysregulation, a dominant clinical risk factor for colonization and invasive disease. As a result, the relative contribution of immune cell abundance versus immune cell functionality in controlling *C. auris* at the skin barrier remains poorly defined.

To address these critical gaps, we developed a physiologically relevant murine model of *C. auris* skin colonization spanning three clinically relevant host conditions: immunocompetent, immunosuppressed and diabetic ketoacidosis (DKA). This approach enables direct comparison of colonization burden, clearance kinetics, local immune responses, and systemic dissemination across distinct immune and metabolic conditions^12,42–44^. By directly contrasting immune suppression with metabolic dysfunction, we revealed significant differences in fungal clearance dynamics, subsequent systemic dissemination and involvement of key immunological factors. This study provides clinically relevant skin colonization models with immunological and metabolic perturbations for mechanistic host-pathogen studies and testing antifungal interventions.

## RESULTS

### Diabetic Ketoacidosis exhibits a high colonization fungal burden and reduced clearance rate

We modeled *C. auris* skin colonization in immunocompetent, immunosuppressed, and streptozotocin (SZT)-induced diabetic-ketoacidosis (DKA) mice (**Figure 1a**). To assess the metabolic and immune status, we measured the blood glucose, urine ketone levels and white blood cell (WBC), respectively (**Supplementary Figure S1a-b**). By day 7 post SZT administration, ∼76% and by day 10, 100% of mice had blood glucose levels >500 mg/dl with urine ketone levels >160 mg/dl, while immunocompetent and immunosuppressed mice had blood glucose levels at 199 ± 32.06 and 196.4 ± 26.87 mg/dl, respectively, with no detectable ketones in the urine (**Supplementary Figure S1a**). Further, immunosuppression resulted into significant reduction in the WBC count compared to both immunocompetent and DKA groups, while DKA mice showed a non-significant decrease in WBC. These trends continued throughout the *C. auris* colonization period staring from day 0 to 20 relative to colonization (**Supplementary Figure S1b**).

**Figure 1.**
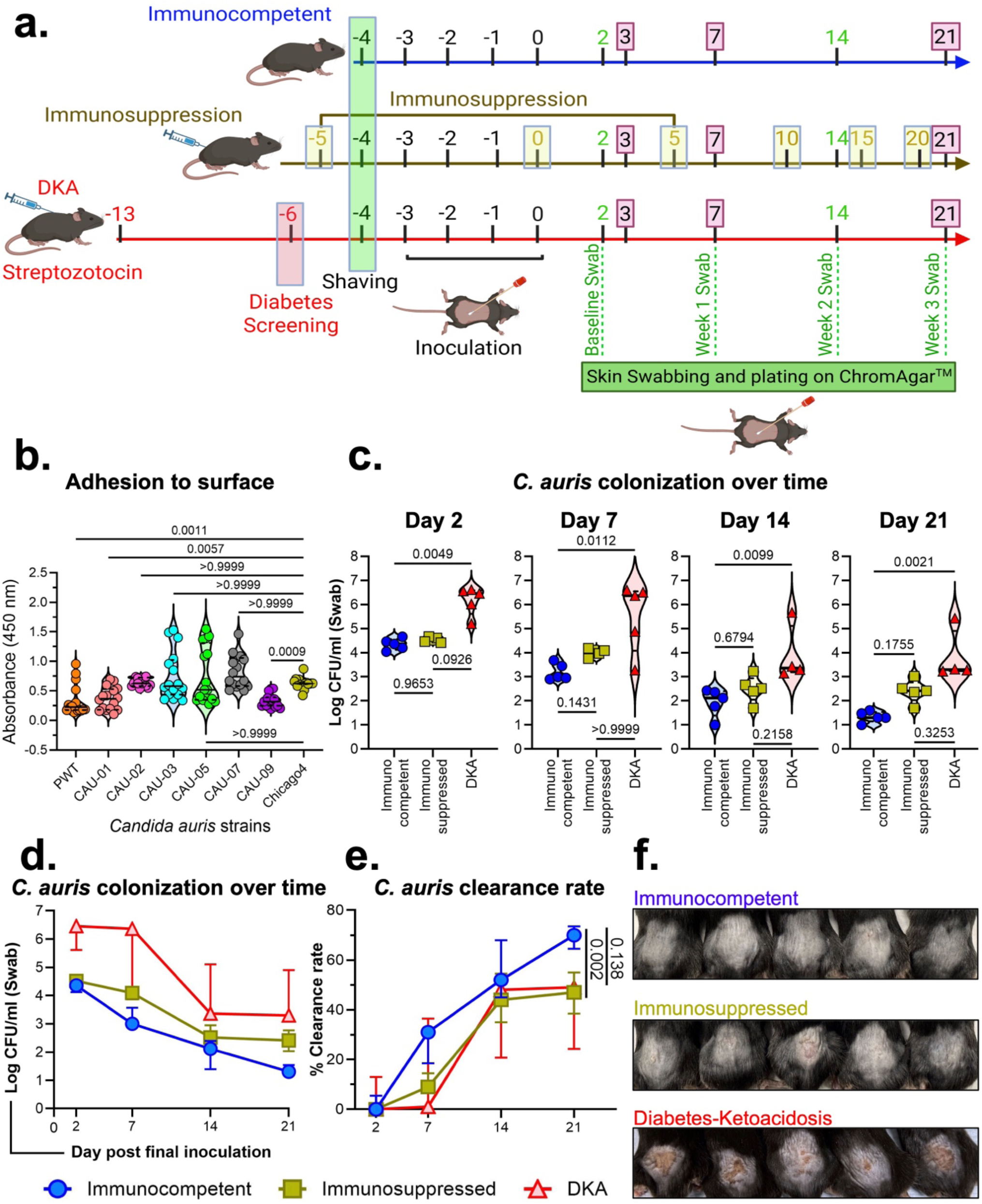
Determination of *C. auris* skin colonization in immunocompetent, immunosuppressed, and diabetic Ketoacidosis (DKA) mice. Immunocompetent, Immunosuppressed and DKA mice (n=5/group) were colonized with 1 × 10^9^ yeast cells/mouse of *C. auris* (Chicago4) for 4 d. At respective time point skin swabs were collected. (**a**) Schematic representation of experimental design. (**b**) Comparative adhesion assessment of different *C. auris* strains to plastic surface. (**c**) CFU per swab from immunocompetent, immunosuppressed, and DKA mice infected with *C. auris* on different days post-colonization is plotted. (**d**) *C. auris* skin colonization monitoring over time compared between the different clinical groups. (**e**) Rate of fungal clearance over time in mice from different clinical groups. (**f**) Visual representation of shaved skin from immunocompetent, immunosuppressed, and DKA mice colonized with *C. auris* after 21 d. All violin plots are expressed as median ± interquartile range. Statistical analysis for all violin plots was performed using the Kruskal-Wallis test followed by Dunn’s multiple comparison test. For statistical significance, a p-value less than 0.05 was considered significant.

We compared several *C. auris* clinical isolates representing clades I-IV for their adhesion ability to substrates. For mouse skin colonization, we prioritized Chicago4 strain which shows higher adhesion to plastic surface compared to PWT, CAU-01, and CAU-09 (p< 0.005), but similar to CAU-02, CAU-03, CAU-05 and CAU-07 (**Figure 1b**), as well as to human keratinocytes^34,61^.

All the mice (back skin) were colonized by day 2 of final inoculation exposure (baseline), which was maintained beyond day 22, as monitored longitudinally by swab sampling on days 7, 14, and 21. Across all time points, DKA mice exhibited significantly higher *C. auris* colonization fungal burden compared to immunocompetent mice (p < 0.05). Specifically, since early time point (day 2, baseline), DKA mice exhibited significantly higher (∼1.42 log-fold, p = 0.0049) skin colonization compared to immunocompetent controls. In contrast, immunosuppressed mice showed a modest, non-significant trend toward a higher (1.04 log-fold, p = 0.9653) burden than immunocompetent controls. Although the initial fungal burden declined by day 14, colonization persisted beyond day 21 across all groups. Throughout the study, DKA mice consistently maintained the highest fungal burden (log 3.296 ± 2.201), followed by immunosuppressed mice (log 2.415 ± 1.322) and immunocompetent mice (log 1.301 ± 0.602) (**Figure 1c**).

There was also a striking difference in the rate at which the mice from different groups cleared the fungus (**Figure 1d, Supplementary Figure S1c**). For instance, post-inoculation, immunocompetent mice achieved 31% clearance from baseline, compared to 9% in immunosuppressed and 1% in DKA mice, indicating a robust early host response in immunocompetent animals. By day 21, immunocompetent mice reached 70% clearance, whereas immunosuppressed and DKA groups plateaued at 47% and 49%, respectively (**Figure 1e**). All these data collectively suggest that a clinical predisposition, such as Diabetic-Ketoacidosis or immunosuppression (neutropenia), facilitates the colonization of *C. auris* on the skin. This was also visibly evident from the scarred skins of DKA mice (**Figure 1f**).

### Diabetic Ketoacidosis results in a high tissue fungal burden and enhanced systemic dissemination

We also determined the skin tissue burden across all three skin colonization models (**Figure 2a**). Interestingly, skin tissue fungal burden in all three mouse models mirrored colonization trends, showing consistent decrease in tissue fungal burden. By day 21 the tissue fungal burden was significantly reduced in all three models (**Figure 2b-d**). Across all time points, DKA mice had significantly higher skin tissue fungal burden compared to immunocompetent mice (**Figure 2e-g**). DKA mice demonstrated the highest fungal load (log 4.771 ± 1.59), which was significantly higher than immunocompetent mice on day 7 (p < 0.0016) and 21 (p < 0.0062). Immunosuppressed mice showed intermediate levels (log 4.146 ± 0.796), which were not significantly different from immunocompetent mice (p > 0.3917) or DKA mice (p > 0.2925). Immunocompetent mice had the lowest tissue burden (log 3.734 ± 0.834) at day 21 (**Figure 2g**). The high tissue fungal burden on the skin of immunosuppressed and DKA mice was also confirmed by histopathology at day 3 and day 7 post-inoculation (**Figure 2h, Supplementary Figure S2**).

**Figure 2.**
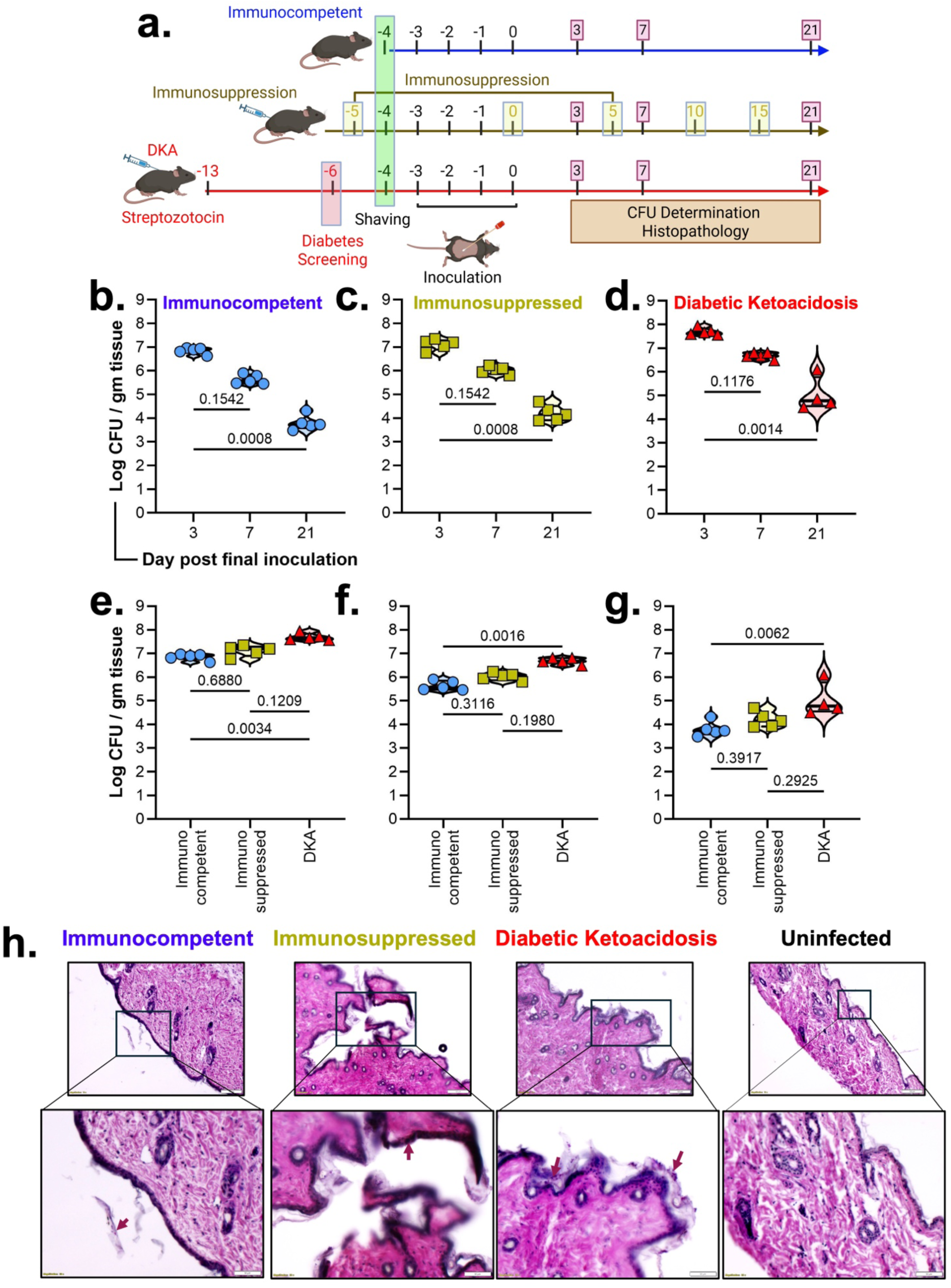
Comparison of Skin tissue fungal burden in immunocompetent, immunosuppressed, and diabetic ketoacidosis mice. Immunocompetent, Immunosuppressed and DKA mice (n=5/group) were colonized with 1 × 10^9^ yeast cells/mouse of *C. auris* (Chicago4) for 4 d. At respective time point skin tissues were collected. (**a**) Schematic representation of experimental design and post-processing of samples. (**b-g**) CFU per g of tissue from immunocompetent, immunosuppressed, and DKA mice infected with *C. auris* on different days post-colonization is plotted. (**b-d**) Within each group, the progress of fungal clearance is plotted as CFU/g of tissue. (**e-g**) CFU per g of tissue is plotted and compared across different clinical groups. The median values with interquartile range are shown. Statistical analysis was performed using the Kruskal-Wallis test followed by Dunn’s multiple comparison test. For statistical significance, a p-value less than 0.05 was considered significant. (**h**) Skin histopathology sections of uninfected and infected mice from different groups (Scale bar: 10 μm) with zoomed insets (Scale bar: 20 μm) are presented.

We also determined the systemic dissemination of *C. auris* in the kidneys at day 21 post-inoculation. Systemic dissemination of *C. auris* was observed in 75% of DKA mice (3/4), whereas no dissemination occurred in immunocompetent or immunosuppressed groups (0%) (**Supplementary Table S4**).

### Immunocompetent, Immunosuppressed, and DKA mice demonstrate differential immune cell infiltration and inflammatory cytokine secretion in *C. auris* colonized skin

*C. auris* skin colonization leads to the invasion of the dermal tissues, which subsequently leads to the inflammation and immune cell infiltration in the skin tissues. While a previous studies demonstrated an infiltration of IL-17A^+^CD4^+^Foxp3^-^, IL-17A^+^γδ and IL-17A^+^CD8^+^ T cells along with IL-17A^+^ILCs at the site of infection ^45^, no studies so far have explored the landscape of infiltrating immune cells at early stage of *C. auris* skin colonization. Primarily, the role of granulocytes including neutrophils and macrophages in the context of *C. auris* skin colonization, remains largely unexplored ^46^. We therefore assessed the infiltration of granulocytes (CD11b^+^), neutrophils (LyG^+^), macrophages (F4/80^+^), dendritic cells (CD11c^+^) and T cells (CD3^+^, CD4^+^) in the *C. auris* colonized skin tissues from our experimental models (**Figure 3a**). The infiltration of granulocyte was significantly reduced (∼20% lower) in both immunosuppressed and DKA mice compared to immunocompetent mice on day 3 post colonization (p < 0.05) (**Figure 3b**). Neutrophil infiltration was also diminished by ∼18% in immunosuppressed mice (p < 0.05) and showed a similar trend in DKA mice (p = 0.089) (**Figure 3c**). By day 7, granulocyte and neutrophil levels increased in DKA mice but remained significantly lower in immunosuppressed mice compared to both immunocompetent controls and DKA mice (p < 0.05) (**Figure 3b-c, Supplementary Figure S3**). Other immune cell subsets (CD3^+^ T cells, F4/80^+^ macrophages, and CD11c^+^ dendritic cells) were comparable among groups at days 3 and 7 post-inoculation (**Figure 3c-d, Supplementary Figure S4**). Interestingly, the percentage of CD4^+^ T_h_ cells was significantly reduced in neutropenic (immunosuppressed) mice compared to immunocompetent mice during the early stage of *C. auris* colonization (**Supplementary Figure S4b,c**), however, during mid-stage colonization, CD4^+^ T_h_ cells started showing up, albeit at a lower percentage, in this group. Collectively, our data suggests that early to mid-stage *C. auris* skin colonization is dependent upon granulocyte infiltration, primarily neutrophils.

**Figure 3.**
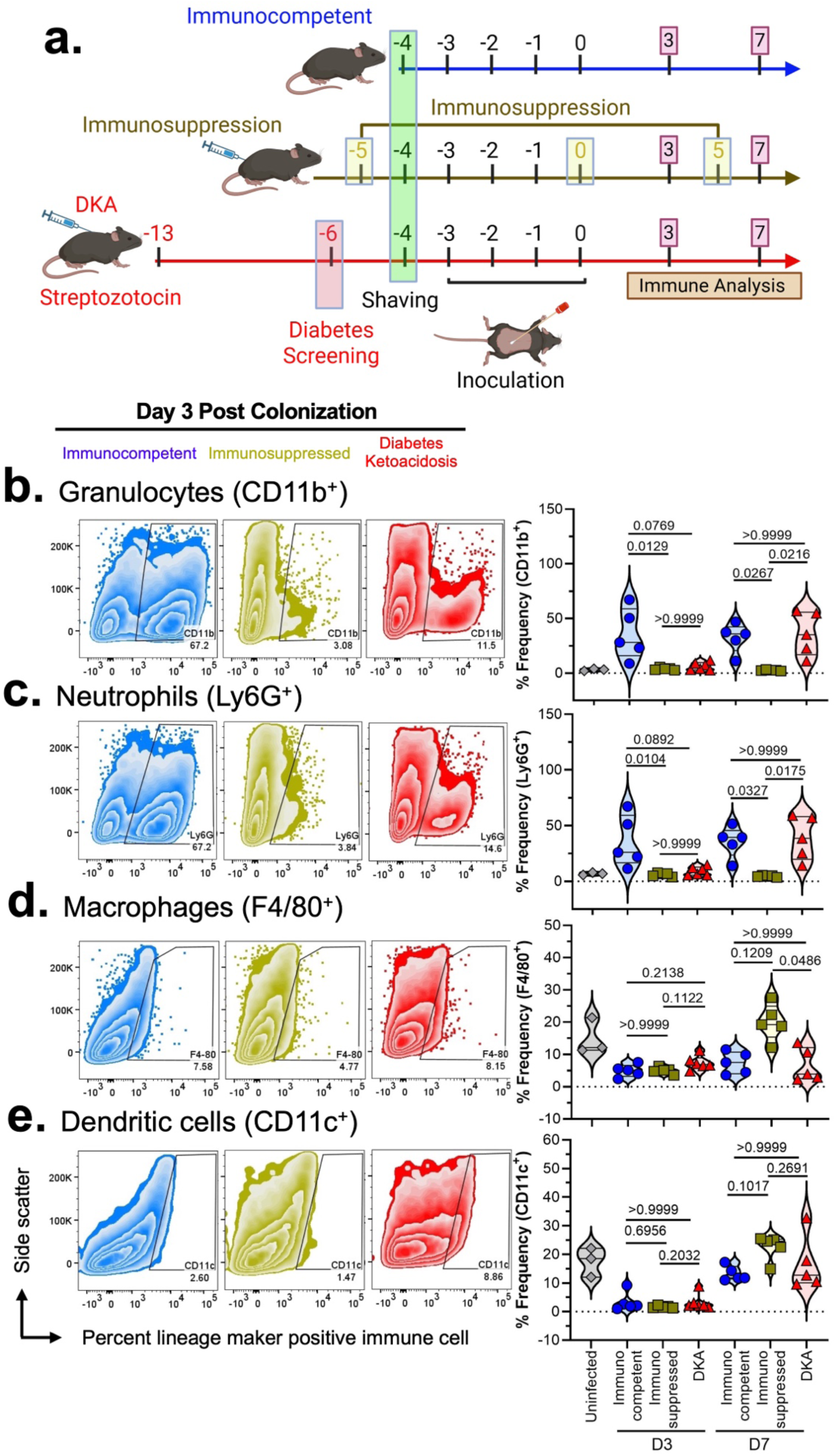
Immunocompetent, Immunosuppressed, and DKA mice demonstrate differential immune cell infiltration in *C. auris* colonized skin. Immunocompetent, Immunosuppressed and DKA mice (n=5/group) were colonized with 1 × 10^9^ yeast cells/mouse of *C. auris* (Chicago4) for 4 d. (**a**) Schematic representation of experimental design and post processing of samples. (**b-e**) 3- and 7-days post colonization, the colonized skin was enzymatically digested and processed to prepare a single-cell suspension and stained with fluorescently labeled antibodies for specific immune cell lineage markers: CD11b for Granulocytes (**b**), Ly6G for Neutrophils (**c**), F4/80 for Macrophages (**d**) and CD11c for dendritic cells (**e**). The stained cells were analyzed using BD FACSymphony, and the area was analyzed with FlowJo v10. Representative scatter graphs and corresponding violin plots (median ± interquartile range) are shown for the different clinical groups. Significant differences in the immune cell populations were analyzed by the Kruskal-Wallis test, followed by Dunn’s multiple comparison test. p values <0.05 were considered statistically significant.

In addition to resident keratinocytes, immune cell infiltration into the skin promotes the formation of a localized inflammatory milieu mediated by secreted cytokines, which can both restrict pathogen growth and dissemination and support tissue repair ^47,48^. We therefore quantified the levels of proinflammatory cytokines and growth factors in the skin of uninfected and *C. auris* colonized mice using cytokine bead array. Skin colonization of *C. auris* led to a robust IL-1α and TNF-α response in immunocompetent mice at both early as well as mid-stage colonization time point (**Supplementary Figure S5a, d**). In contrast, cytokine responses were markedly reduced in immunosuppressed mice, while DKA mice showed no significant reduction. IL-1β was minimally detected in the skin following *C. auris* colonization (**Supplementary Figure S5b**), and MCP-1 levels remain unchanged across groups at the early time point (**Supplementary Figure S5c**). Other proinflammatory cytokines including, IL-6, TGFβ, IL-23, GM-CSF, IL-12p70, IFNγ and IL-27 were undetectable in the skin at both early as well as mid-stage infection.

Levels of the anti-inflammatory cytokine, IL-10, were not significantly different among different groups at mid stage of colonization (**Supplementary Figure S5e**). A hallmark of *C. auris* skin infection is the infiltration of T_h_17 and γδT-cells creating an IL-17A/IL-17F/IL-22 environment. However, this infiltration occurs at a later stage. Therefore, it was not surprising that in our model, IL-17A was not detected in either group at both early and mid-stage of *C. auris* colonization, except in DKA mice, which showed significantly higher levels of Il-17A at mid-stage of colonization (**Supplementary Figure S5f**).

### Bone marrow neutrophils and differentiated macrophages from DKA mice exhibit impaired anti-fungal responses

Innate immune cells such as neutrophils and macrophages are the first line of defense against any invading pathogens. Our data also supports the hypothesis that neutrophils are one of the first responders during *C. auris* skin colonization (**Figure 3B**). While infiltrating neutrophil numbers are significantly reduced in DKA mouse skins during the early phase of colonization, their numbers increase considerably by day 7 post colonization. In fact, when we profiled the bone marrow, we observed that neutrophil numbers were significantly elevated in DKA mice bone marrow compared to immunocompetent mice (**Supplementary Figure S6a**). Interestingly, resident macrophage numbers were not altered in the bone marrow of DKA mice compared to immunocompetent mice (**Supplementary Figure S6b**), mimicking an observation similar to the infected skin microenvironment. Despite this increase in neutrophil numbers, the overall fungal burden in the DKA mouse skin remains significantly higher, suggesting that despite their presence, an impaired antifungal response exists in these neutrophils. We, therefore, next characterized the neutrophil antifungal response, infecting bone marrow neutrophils from immunocompetent and DKA mice with *C. auris ex vivo* (**Figure 4a**). As predicted, neutrophils from DKA mouse bone marrow phagocytosed significantly less *C. auris* compared to the same from immunocompetent mice (**Figure 4b, c**). Phagocytic uptake of pathogens is often associated with the generation of NADPH-dependent ROS, a ploy by innate immune cells to eradicate the invading pathogen^49^. We therefore investigated if *C. auris*-dependent ROS was also affected in DKA neutrophils. Interestingly, we observed an opposing trend, wherein DKA neutrophils generated significantly more ROS upon *C. auris* invasion, compared to immunocompetent neutrophils (**Figure 4d, e**). While it would be expected that a higher ROS response would lead to efficient fungal killing, the same was not observed with DKA neutrophils. DKA neutrophils were approximately 20% less efficient in killing *C. auris* compared to immunocompetent neutrophils (Median 42% in Immunocompetent vs median 24% DKA) (**Figure 4f**). However, compared to immunocompetent neutrophils, DKA neutrophils were approximately 10% more sensitive to *C. auris*-mediated cell death (**Figure 4g**).

**Figure 4.**
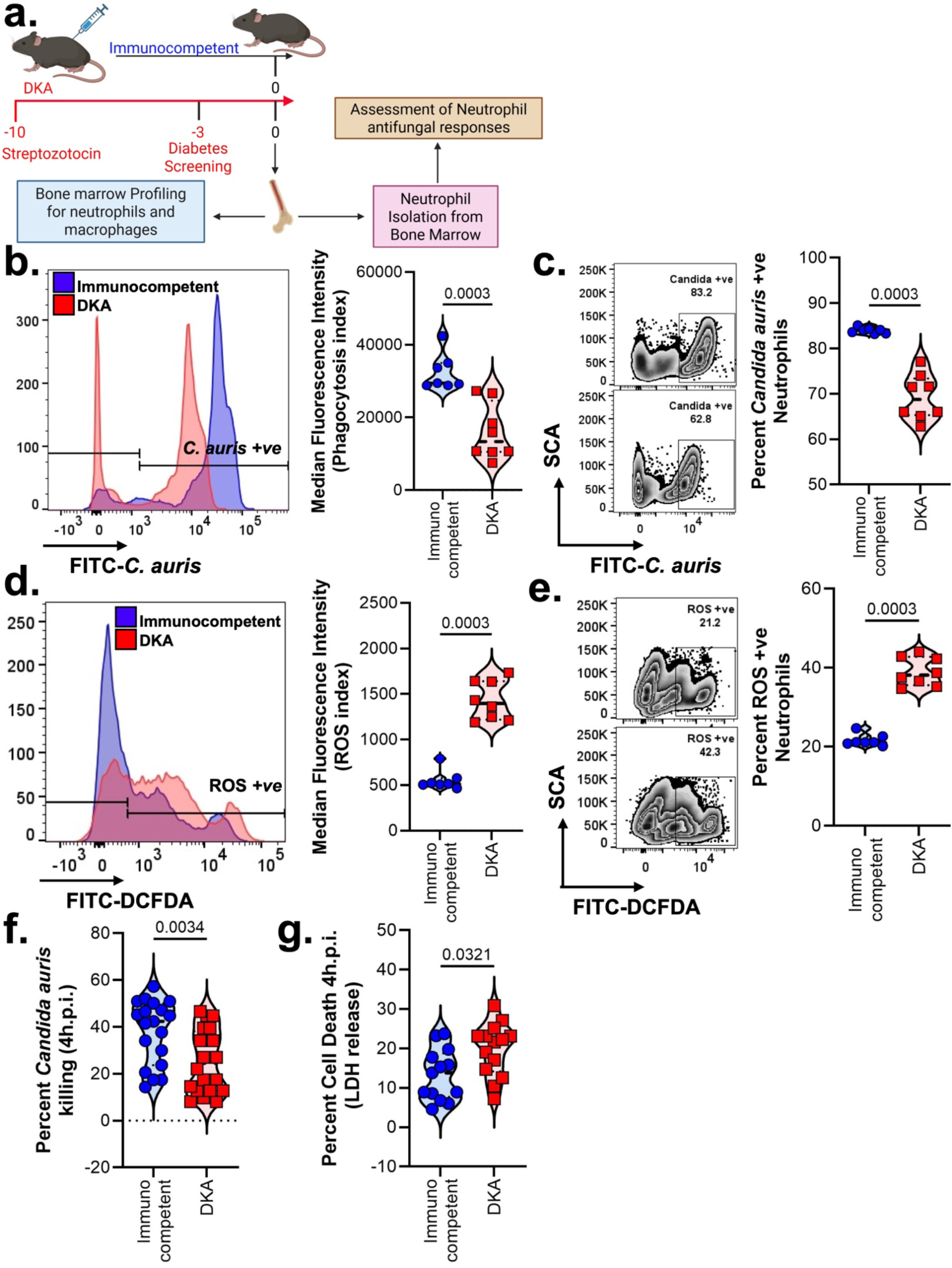
Characterizing the antifungal response of neutrophils from DKA mice. Bone marrow neutrophils from immunocompetent and DKA mice were isolated and antifungal responses were characterized. (**a**) Schematic representation of experimental design. (**b-c**) Neutrophils were infected with CFSE-labelled *C. auris* (MOI-1) for 1.5 h. Phagocytosed fungus was determined by flow cytometry. Representative histogram plot and calculated median fluorescence intensity are presented (**b**). Representative zebra plot and calculated percentage *C. auris* positive neutrophils are presented (**c**). (**d-e**) Neutrophils were infected with *C. auris* (MOI-1) for 1 h. Total cellular ROS was determined by flow cytometry. Representative histogram plot and calculated median fluorescence intensity are presented (**d**). Representative zebra plot and calculated percentage ROS positive neutrophils are presented (**e**). (**f-g**) Neutrophils were infected with *C. auris* (MOI-5) for 4 h. Percentage killing of *C. auris* was determined by the ratio of CFU of *C. auris* surviving in the presence or absence of neutrophils (**f**). Percentage neutrophil death upon *C. auris* infection was determined by assessing the amount of LDH released (**g**). All violin plots are expressed as median ± interquartile range. Statistical analysis was performed using the two-tailed Mann Whitney test. For statistical significance, a p-value less than 0.05 was considered significant.

Bone marrow cells from immunocompetent and DKA mice were differentiated into primary macrophages using GM-CSF *in vitro*. Despite being in the absence of any hyperglycemic and ketogenic stimuli, differentiated macrophages from DKA bone marrow exhibited similar impaired *C. auris* phagocytic uptake vs macrophages from immunocompetent mice (**Figure 5a, b**). However, contrary to neutrophils from DKA mice, DKA macrophages did not generate a higher *C. auris*-dependent ROS (**Figure 5c, d**). The overall ability of macrophages to eliminate *C. auris* was significantly impacted in DKA macrophages, similar to neutrophils (**Figure 5e**). Interestingly, unlike neutrophils, DKA macrophages were resistant to *C. auris*-mediated cell death (**Figure 5f**). Hyperglycemia is often associated with elevated levels of proinflammatory cytokines secreted by macrophages ^50,51^. We therefore profiled secreted levels of proinflammatory cytokines released by primary macrophages from DKA and immunocompetent mice upon acute *C. auris* infection. As expected, secreted levels of IL-1α, IL-1β, IL-12p70 and IFNβ were significantly elevated upon *C. auris* exposure to primary macrophages from DKA mice compared to their immunocompetent counterparts (**Figure 5g, Supplementary Figure S7**). Although not significant, IL-6 and TNF-α also showed a similar trend. Interestingly, secreted levels of the anti-inflammatory cytokine, IL-10, were also elevated in these macrophages. The macrophage-specific chemokine, MCP-1, was, however, modestly but significantly altered between the different groups. These findings are not similar to the overall cytokine landscape observed in the infected skin. However, it is worth noting that the skin cytokine microenvironment is contributed not only by infiltrating immune cells but also by resident keratinocytes. How keratinocytes in DKA mice respond to *C. auris* infection needs further investigation.

**Figure 5.**
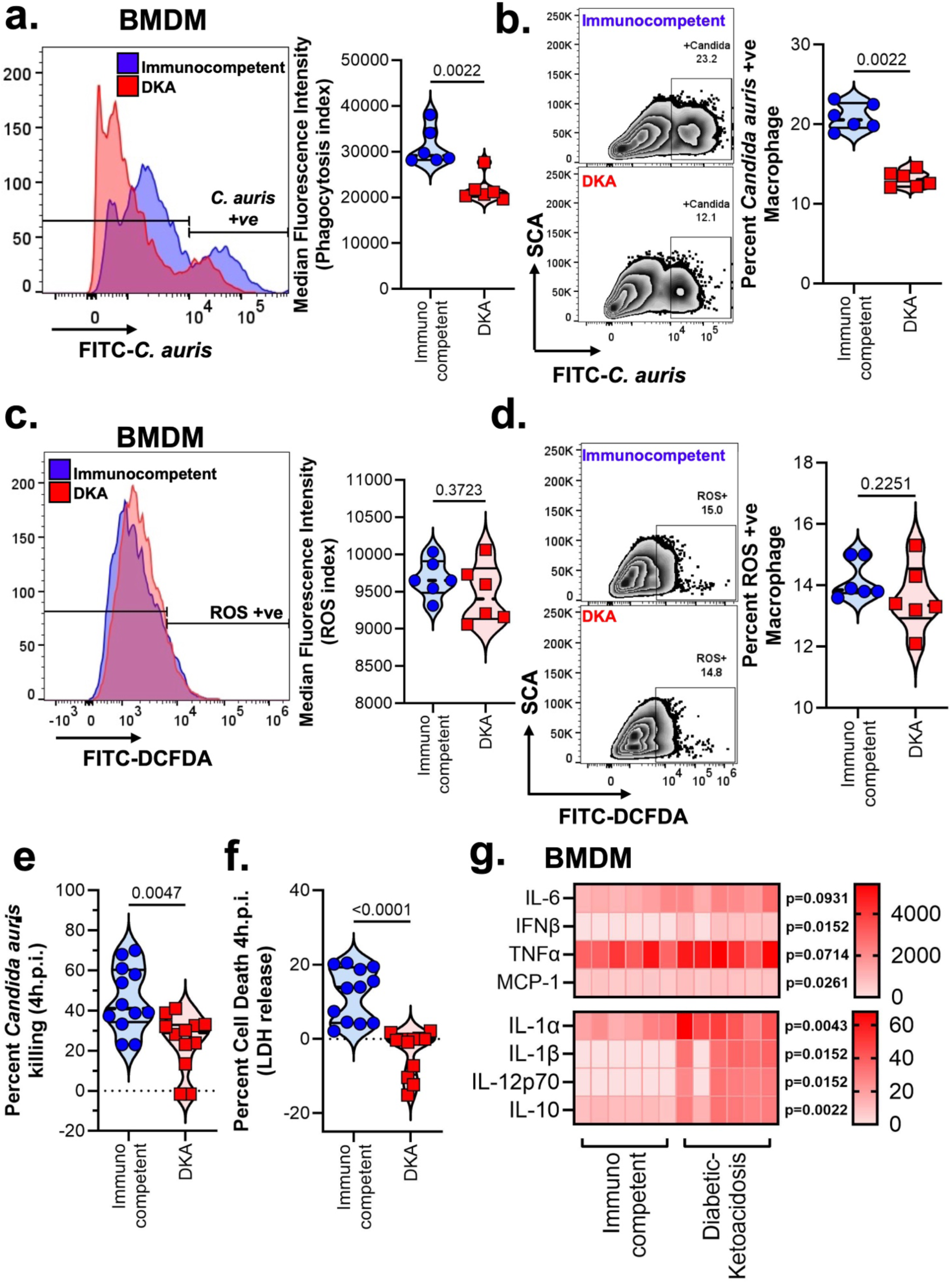
Characterizing the antifungal response of primary macrophages from DKA mice. Primary macrophages from Bone marrow cells of immunocompetent and DKA mice, were differentiated using GM-CSF. (**a-b**) Macrophages were infected with CFSE-labelled *C. auris* (MOI-1) for 1.5 h. Phagocytosed fungus was determined by flow cytometry. Representative histogram plot and calculated median fluorescence intensity are presented (**a**). Representative zebra plot and calculated percentage *C. auris* positive neutrophils are presented (**b**). (**c-d**) Macrophages were infected with *C. auris* (MOI-1) for 1 h. Total cellular ROS was determined by flow cytometry. Representative histogram plot and calculated median fluorescence intensity are presented (**c**). Representative zebra plot and calculated percentage ROS positive neutrophils are presented (**d**). (**e-g**) Macrophages were infected with *C. auris* (MOI-5) for 4 h. Percentage killing of *C. auris* was determined by the ratio of CFU of *C. auris* surviving in the presence or absence of macrophages (**e**). Percentage neutrophil death upon *C. auris* infection was determined by assessing the amount of LDH released (**f**). Cell culture supernatants were analyzed for levels of secreted cytokines by cytokine bead-array. Cytokine concentrations are expressed as a heatmap (**g**). All violin plots are expressed as median ± interquartile range. Statistical analysis for all data was performed using the two-tailed Mann Whitney test. For statistical significance, a p-value less than 0.05 was considered significant.

### DKA conditions are associated with decreased expression of immune receptors, proteases and an impaired antifungal response in neutrophils and macrophages

To get a mechanistic insight into why innate immune cells from DKA mice exhibited an impaired antifungal response compared to the same from immunocompetent mice, we performed gene expression analysis, focusing on surface recognition receptors and antimicrobial proteases. *C. auris* infection induced the expression of both *Tlr2* as well as *Clec7a* (Dectin-1) in neutrophils from immunocompetent mice, receptors involved in the recognition of fungal cell wall ^52^ (**Figure 6a**). While the expression of *Tlr2* was not altered in DKA neutrophils, *Clec7a* mRNA levels were significantly reduced in DKA neutrophils upon *C. auris* infection. The gene expression of the neutrophil specific enzyme, Myeloperoxidase, was not altered upon *C. auris* infection, but the basal gene expression levels as well as levels upon *C. auris* infection, was significantly downregulated in DKA neutrophils compared to immunocompetent neutrophils (**Figure 6a**). A similar significant reduction was also observed for genes encoding serine proteases including Elastase, Cathepsin G and Proteinase 3 in DKA neutrophils. Serine proteases play a crucial role in antifungal defense ^53^ and a reduction in the gene expression of these proteases possibly explains why neutrophils from DKA mice exhibit an impaired antifungal response. Consistent with the gene expression data, neutrophil elastase secretion in the supernatant post *C. auris* infection was significantly increased in the neutrophils from immunocompetent mice, compared to DKA mice (**Figure 6b**). A similar trend was also observed in the total cell lysates of these neutrophils, further confirming that hyperglycemia and ketoacidosis contribute to an impaired antifungal response via a defective NETosis response. We also profiled selective zinc-dependent metalloproteinases that not only play a role in tissue remodeling and wound healing but also contribute to protection from invading pathogens^54^. Of note, MMP7, MMP8, MMP9, MMP19 and MMP28 have been implicated inflammatory responses and closely associated infections. We observed that in DKA neutrophils the mRNA expression of *Mmp7*, *Mmp8* and *Mmp9* were strongly downregulated compared to immunocompetent neutrophils, while *Mmp19* and *Mmp28* were largely unaffected (**Figure 6a**).

**Figure 6:**
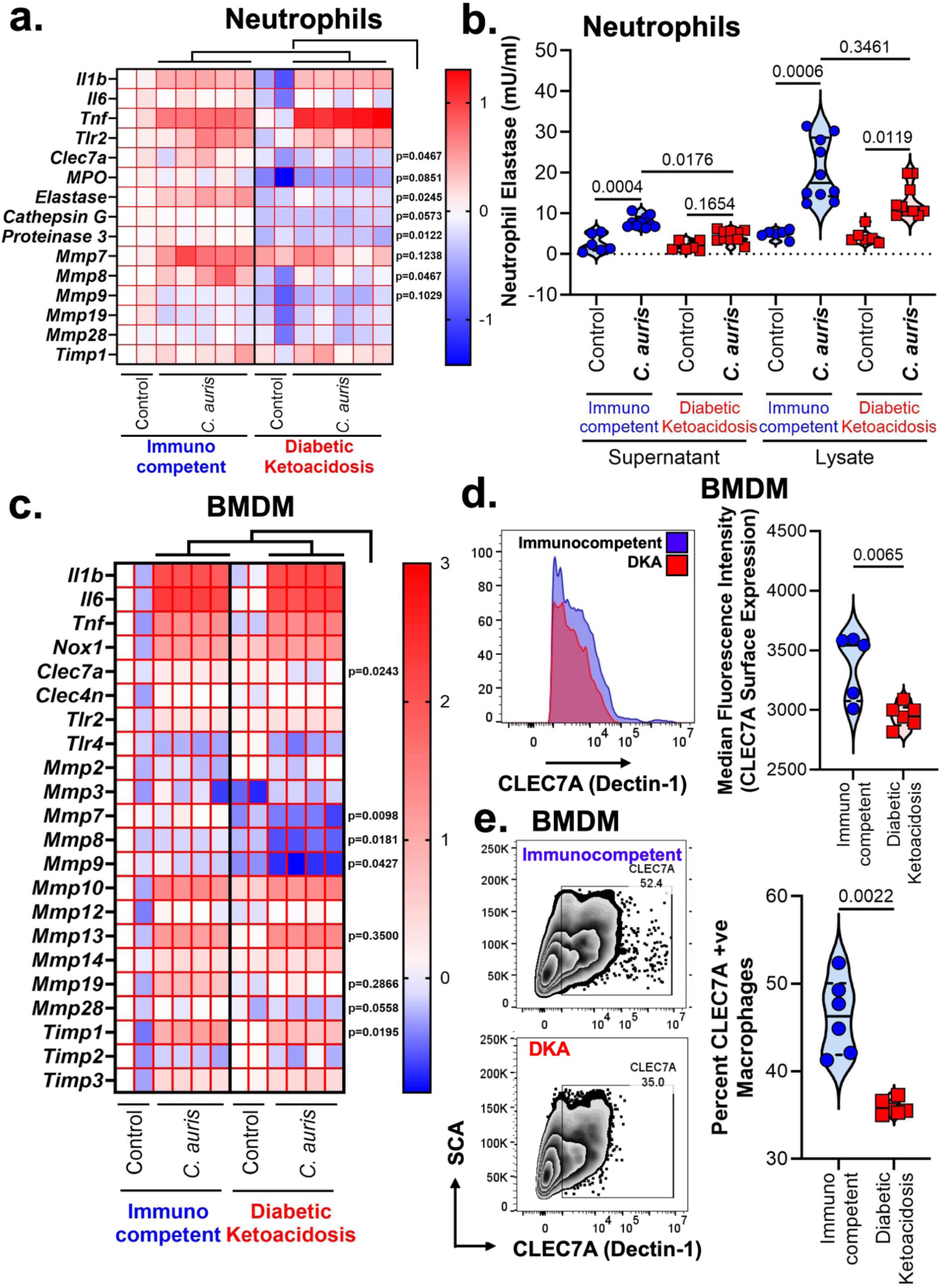
Expression of surface receptors and proteases involved in inflammatory and antimicrobial responses are altered in DKA neutrophils and macrophages. (**a**) Bone marrow Neutrophils from immunocompetent and DKA mice were infected with *C. auris* (MOI-5) for 4 h. Total RNA was isolated and expression of mRNA for the listed genes was assessed using qPCR. Gene expressions are expressed as a heatmap. (**b**) Neutrophil elastase levels, secreted in the supernatant and present in the cell lysate, in the presence or absence of *C. auris* was determined. (**c**) Bone marrow derived macrophages from immunocompetent and DKA mice were infected with *C. auris* (MOI-5) for 4 h. Total RNA was isolated and expression of mRNA for the listed genes was assessed using qPCR. Gene expressions are expressed as a heatmap. (**d-e**) Surface expression of CLEC7A (Dectin-1) in primary macrophages was determined by flow cytometry. Representative histogram plot and calculated median fluorescence intensity are presented (**d**). Representative zebra plot and calculated percentage *C. auris* positive neutrophils are presented (**e**). All violin plots are expressed as median ± interquartile range. Statistical analysis for (**a-c**) was performed using Kruskal-Wallis test followed by Dunn’s multiple comparison test while the same for (**d-e)** was performed using the two-tailed Mann Whitney test. For statistical significance, a p-value less than 0.05 was considered significant.

Primary macrophages from DKA mice also showed a similar phenotype. Notably, the expressions of *Clec7a*, *Mmp7*, *Mmp8, Mmp9* were significantly reduced upon *C. auris* infection in DKA primary macrophages (**Figure 6c**). *Mmp13*, *Mmp19* and *Mmp28* also showed similar trends. The inhibitor of matrix metalloproteinases (TIMP) also plays an important role in tissue healing with TIMP1 additionally sustaining myeloid-biased hematopoiesis after polymicrobial peritonitis ^55^. Interestingly, *Timp1* was strongly induced upon *C. auris* infection, however this induction was significantly less in DKA macrophages, compared to immunocompetent macrophages (**Figure 6c**). Other genes including various MMPs and TIMPs were not significantly altered in DKA macrophages. Lastly, we validated the surface expression of receptors involved in fungal cell wall recognition by flow cytometry. As expected, DKA macrophages expressed significantly lesser Dectin-1 (CLEC7A) (**Figure 6d, e**) compared to their immunocompetent counterparts. Interestingly, while the total expression of surface TLR2 was significantly higher in DKA macrophages (**Supplementary Figure S8b**), the number of receptors per cell (assessed by median fluorescence intensity) was higher in immunocompetent macrophages (**Supplementary Figure S8a**). The overall expression of TLR4 remained unchanged in between the two groups (**Supplementary Figure S8c, d**). Collectively, our data suggest a unique role of serine proteases, matrix metalloproteinases and surface receptors in hyperglycemic and ketoacidosis conditions, working in conjunction to impart an effective antifungal response in neutrophils and macrophages.

### Hyperglycemia and Ketoacidosis modulate *C. auris* cell wall architecture to influence host inflammatory and antimicrobial responses

It is widely appreciated that the complex cell wall architecture of *C. auris* influences host inflammatory and antimicrobial responses ^56,57^. The cell wall of a fungus is a dynamic entity that is remodeled under environmental stress, thereby influencing its virulence ^59^. We therefore hypothesized that hyperglycemia and ketoacidosis under DKA conditions modulate the cell wall architecture of *C. auris*, thereby promoting skin colonization and virulence. We observed that the growth of Chicago4 was significantly enhanced in the presence of hyperglycemic condition (≥4 mg/ml glucose) but not under ketogenic (β-hydroxy butyrate [BHB], ≥ 10 mM) conditions. Further, a combination of both hyperglycemic and ketogenic conditions did not further enhanced *C. auris* growth vs hyperglycemic condition alone (**Figure 7a**). To ensure that this was not a clade specific phenomenon, we tested the growth of the highly virulent, Clade I *C. auris* (CAU-09) under similar conditions. As expected, CAU-09 showed similar growth pattern as Chicago4 (**Supplementary Figure S9a**). We next profiled the cell wall architecture of these strains by flow cytometry. Interestingly, a combination of hyperglycemic (high glucose) and ketonic (high BHB) conditions significantly reduced the mannan content of both Chicago4 and CAU-09 (**Figure 7b, Supplementary Figure S9b**), whilst alone failed to reduce it significantly. In contrast, cell wall chitin levels trended higher in a combination of hyperglycemic and ketonic conditions vs normal control condition (**Figure 7c, Supplementary Figure S9c**). In addition to polysaccharides, adhesin proteins constitute an important component of *C. auris* cell wall, not only aiding in adhesion to host cells but also contributing to virulence ^34,37^. Interestingly, changes in local environment also modulate the expression of these adhesin proteins^63^. We therefore profiled the expression of known adhesin proteins from *C. auris*. The Als ortholog *PIS50263.1* was significantly upregulated upon exposure to both hyperglycemic and ketonic condition alone or in combination, with the other Als orthologs showing similar trend (**Figure 7d**). Amongst the different members of the HYR family, these conditions downregulated the expression of genes encoding HIL-2, HIL-3, HIL-4 and HIL-6, but showed upregulation HIL-7, and HIL-8 showing a similar trend, suggesting that amongst the different members of HIL family genes. To further validate this finding, Chicago4, grown under the hyperglycemic or ketonic condition or in both, were incubated with anti-Als3p (generated through NDV-3A vaccination)^33^, anti-Hyr1^35,64^ or control mouse sera. We have previously confirmed that the mouse sera containing anti-Als3 or anti-Hyr1 antibodies cross-react with *C. auris* Als orthologs and HIL family proteins. Consistent with the gene expression data, anti-Als3p and anti-Hyr1p sera showed enhanced antibody binding with Chicago4, when it was grown in under hyperglycemic but not under ketonic condition (**Figure 7e**). This observation was further replicated using the clade I strain, CAU-09 (**Supplementary Figure S9d**).

**Figure 7:**
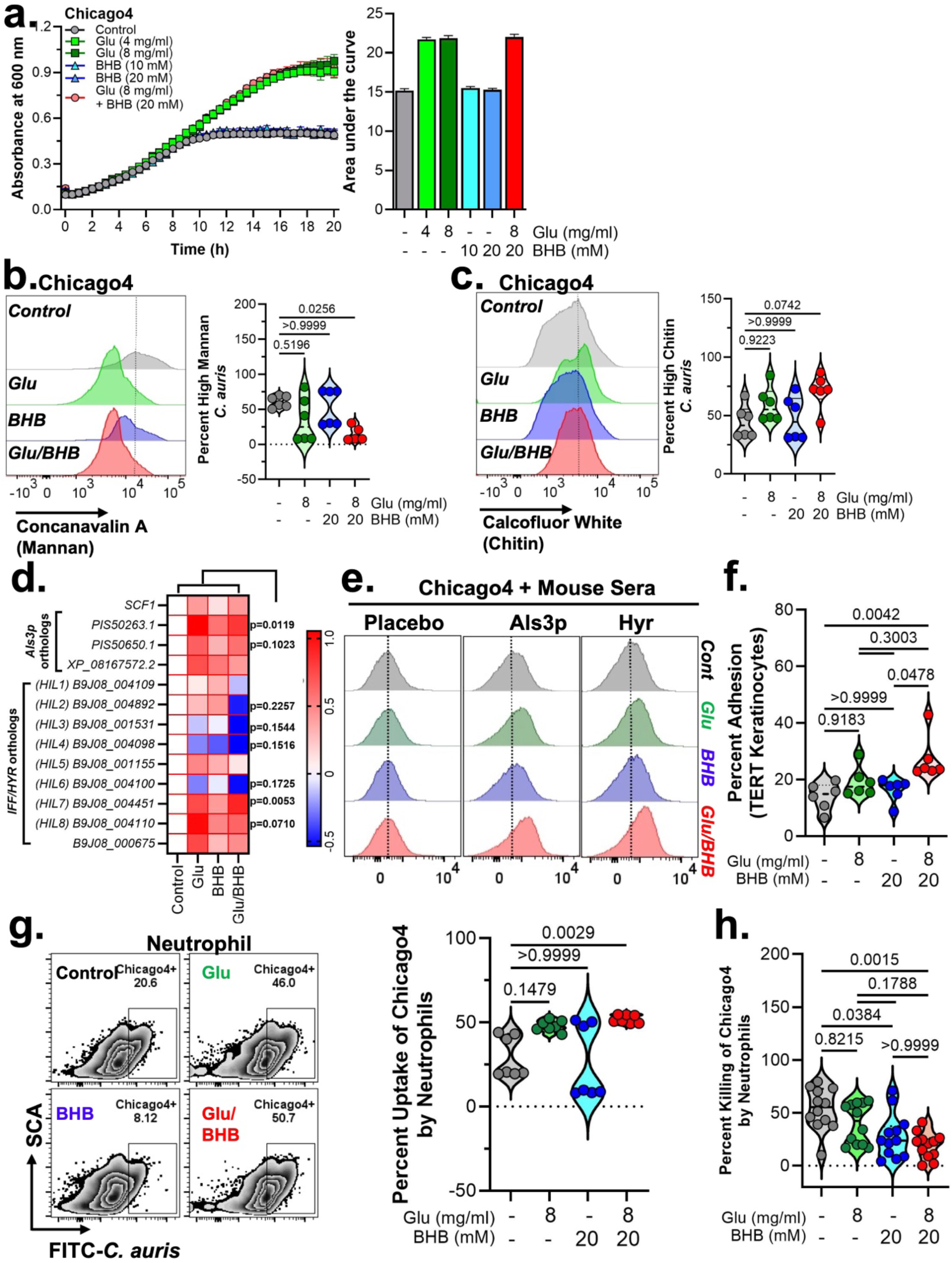
Hyperglycemia and ketone bodies modulate cell wall components of *C. auris* influencing its skin colonization and virulence. (**a**) The clade IV strain Chicago4 was grown in the presence or absence of glucose or BHB alone or in combination for 20 h. Absorbance was measured at 600 nm as a measure of growth. Area under the curve was calculated thereafter. (**b-d)** Chicago4 was grown in the presence or absence of glucose or BHB alone or in combination overnight. Fungal cells were fixed and cell wall was stained for mannan (**b**) and chitin (**c**). Expression of these cell wall components were then analyzed by flow cytometry. Representative histogram plot and calculated percentage high cell wall components are presented. (**d**) Total RNA was isolated from Chicago4 grown under the high glucose and BHB conditions and expression of mRNA for the listed genes was assessed using qPCR. Gene expressions are expressed as a heatmap. (**e**) Fixed Chicago4 cells grown under the high glucose and BHB conditions were incubated with either control, anti-Als3p or Anti-Hyr1p mouse sera. Antibody binding to fungal cell wall was detected with Mouse IgG antibodies conjugated with Alexa Fluor 488 using flow cytometry. Representative histogram plots are presented. (**f**) Adhesion of Chicago4, grown under different conditions, to N/TERT-1 keratinocytes was determined by CFU method. (**g-h**) Uptake of Chicago4 (**g**) and percentage killing of the fungus (**h**), grown under different conditions, by neutrophils was calculated by flow cytometry and CFU method respectively. All violin plots are expressed as median ± interquartile range. Statistical analysis for (**b-d, f-h**) was performed using Kruskal-Wallis test followed by Dunn’s multiple comparison test while the same for (**d)** was performed using two-way ANOVA followed by Dunnett’s post hoc test. For statistical significance, a p-value less than 0.05 was considered significant.

Further, we investigated whether the enhanced expression of adhesins would result in increased adhesion to host cells. As expected, adhesion of Chicago4 to N/TERT-1 keratinocytes^61^ was significantly increased when the fungus was grown in the presence of combination of hyperglycemic and ketonic conditions but did not show increase when grown in each condition alone (**Figure 7f**). Since, modulated cell wall architecture is associated with immune modulation, particularly innate cell responses. Next, we investigated with the change in cell wall polysaccharides and adhesins under hyperglycemic and ketonic conditions leads to neutrophil and macrophage uptake and killing. Indeed, Chicago4 by exposure to a combination of glycemic and ketonic conditions leads to significantly increased uptake by neutrophils and primary macrophages, and exposure to ketonic condition alone only increased Chicago4 uptake by macrophages (**Figure 7g, Supplementary Figure S10a**). However, despite enhanced uptake ketonic condition alone or in combination with hyperglycemic condition leads to significantly decreased in fungal killing by neutrophils indicating dysfunctional killing activity (**Figure 7h**).

Next, we investigated if the dysfunctional antifungal activity of neutrophils and macrophage is associated with altered immune signatures. Indeed, Chicago4 exposed to both hyperglycemic and ketonic conditions leads to significantly higher IL-1β response in macrophages and neutrophils (**Supplementary Figure S10b**), compared to unexposed fungus or fungus exposed to either condition alone. The release of TNF-α from macrophages and neutrophils was significantly enhanced, while IL-6 levels remained unchanged (**Supplementary Figure S10c, d**).

## DISCUSSION

*C. auris* is a multidrug-resistant fungal pathogen with an exceptional capacity for persistent skin colonization and nosocomial transmission, presenting a major challenge for infection prevention in healthcare settings ^3,27,58,60,62,65^. Clinical surveillance consistently links colonization and invasive diseases to host immune and metabolic dysfunction, however, the mechanism by which these states shape colonization burden, persistence, and dissemination risk remain poorly defined ^26,58,66–68^. Here, we establish complementary murine model of *C. auris* skin colonization across immunocompetent, immunosuppressed, and diabetic ketoacidosis (DKA) conditions, enabling direct dissection of how immune suppression versus metabolic dysregulation differentially remodel cutaneous host defense. This study reveals host metabolic dysfunction, not immune cell scarcity, as primary driver of persistent *C. auris* skin colonization and dissemination, identifying qualitative defects in innate antifungal immunity as a key determinant of invasive risk.

Consistent with previous reports^44^, immunosuppressed mice failed to restrict *C. auris* skin colonization, likely reflecting quantitative deficiencies in circulating and tissue-resident neutrophils. In contrast, SZT-induced DKA further exacerbated fungal persistence despite preserving granulocyte and neutrophil recruitment to the skin. This dissociation between immune cell abundance and fungal control indicates that diabetes imposes a qualitative defect in neutrophil antifungal function rather than a numerical deficit. Supporting this interpretation, prior work demonstrates that SZT-induced diabetes disrupts multiple neutrophil effector pathway, including impaired phagocytosis, defective autophagy, increased DNA damage, and dysregulated reactive oxygen species (ROS) responses following activation^69–71^. In parallel, reduced macrophage inflammatory protein-2 (MIP-2) expression reported in diabetic macrophages suggests impaired macrophage-neutrophil coordination, further constraining effective innate immune responses. Together, these findings provide a mechanistic framework in which metabolic dysregulation uncouples neutrophil recruitment from antifungal efficacy, explaining the failure of DKA mice to restrict *C. auris* colonization despite preserved immune cell infiltration.

While late-stage host responses to *C. auris* skin colonization have been characterized by T_h_17 and IFNγ-driven adaptive immunity^72^ ^73^, little is known about the early innate immune events that determine colonization outcomes. Our data highlights a dominant role for granulocytes, particularly neutrophils, during early and mid-stage colonization, with impaired early infiltration correlating with higher fungal burden. Notably, although neutrophil numbers recover in DKA skin at later time points, this quantitative rebound fails to translate into effective fungal control, reinforcing the concept that neutrophil functionality, rather than abundance alone, dictates colonization outcomes under metabolic stress.

Cytokine profiling of colonized skin revealed surprisingly modest inflammatory responses during early and mid-stage colonization, in contrast to robust cytokine production observed in *ex vivo* macrophage assay. This divergence likely reflects the unique immunological constraints of the skin microenvironment, where resident keratinocytes, stromal cells, and infiltrating immune populations collectively shape local inflammatory tone. While pro-inflammatory cytokines, such as TNF- α, and IL-1 family members are known to contribute to fungal containment and tissue repair^74^, their limited induction during early colonization suggests that *C. auris* may initially evade or suppress cutaneous inflammatory signaling. The delayed emergence of IL-17A exclusively in DKA skin further supports the notion that adaptive immune engagement is a secondary event, occurring after prolonged fungal persistence rather than directly mediating early containment.

Mechanistically, our data identify coordinated defects in pattern-recognition receptor, serine proteases, and matrix metalloproteinases (MMPs) as central contributors to impaired antifungal immunity under DKA conditions. Reduced expression of CLEC7A (Dectin-1), neutrophil elastase, and antifungal proteases, despite preserved or elevated ROS generation, suggest a decoupling of fungal recognition, killing and tissue remodeling programs. Similarly, diminished expression of MMPs and TIMP1 in DKA phagocytes points to broader defects in inflammatory regulation and would repair, processes increasingly recognized as integral to host defense at epithelial barriers^75,76^. Collectively, these findings position metabolic dysregulation as a state that rewires innate immune effectors pathways, rendering phagocytes numerically present yet functionally effective.

We further demonstrate that the combined metabolic stresses characteristic of hyperglycemia and ketoacidosis selectively reprogram *C. auris* cell wall architecture and adhesin expression, with direct consequences for host innate immune interactions. The observed reduction in surface mannan together with increased chitin exposure under combined glucose and ketone conditions suggests a coordinated remodeling response that may alter pathogen-associated molecular pattern (PAMP) presentation and immune recognition. Such dynamic restructuring of the fungal cell wall is consistent with prior evidence that *C. auris* modulates its surface architecture in response to environmental cues to influence host-pathogen interactions and virulence. Importantly, our finding that adhesin expression, particularly ALS orthologs and select HIL family members, is enhanced under these metabolic conditions supports a model in which nutrient availability directly drives fungal adhesion capacity and epithelial colonization potential. This is in line with recent work showing that *C. auris* uses nutrient sensing to fine-tune virulence traits, including adhesion and immune engagement, depending on local environmental inputs ^63^.

Functionally, these structural changes translate into increased keratinocyte adhesion and enhanced recognition by phagocytes yet paradoxically result in impaired neutrophil killing despite increased uptake. This uncoupling of phagocytosis from fungicidal activity suggests that metabolic conditioning of *C. auris* promotes a state of immune activation without effective clearance, a phenotype reminiscent of reports that *C. auris* can persist within phagocytes and resist intracellular killing mechanisms. The concomitant increase in IL-1β and TNF-α production further supports the notion that remodeled cell wall components and adhesins amplify inflammatory signaling, likely through altered engagement of pattern recognition receptors. Clinically, these findings are highly relevant in the context of diabetes and DKA, where hyperglycemia and altered, metabolic states are known to impair neutrophil antimicrobial function and predispose patients to fungal infections. Thus, our data suggest that DKA-associated metabolic cues not only weaken host antifungal defenses but also actively reprogram *C. auris* into a more adhesive and immunomodulatory state, thereby promoting persistence at the skin interface ^82,84^.

Clinically, diabetes is a well-established risk factor for *C. auris* persistent colonization, and progression to invasive diseases^26,65–68,77–79^. Our findings provide mechanistic insight into these epidemiological associations, demonstrating that DKA uniquely permits systemic dissemination from cutaneous sites despite intact immune cell recruitment, whilst also modulating the cell wall architecture of *C. auris,* which subsequently induced altered antifungal responses. These data have direct implications for infection prevention strategies, supporting enhanced surveillance, prolonged isolation, and targeted decolonization efforts in metabolically high-risk patients. Moreover, identifying functional deficits, rather than simple leukocyte counts, as predictors of dissemination risk may inform future biomarker development.

Although this study identified several key factors that dictates the *C. auris* skin colonization, yet several limitations exist. The SZT-induced DKA model captures key features of acute metabolic dysregulation but does not fully recapitulate the heterogeneity of diabetes in human patients. Additionally, our analyses focused primarily on innate immune compartments, defining how keratinocytes and adaptive immune sub-sets integrate with these pathways will be important area for future investigation. Despite these limitations, our model provides a robust platform for disentangling immune versus metabolic contributions to fungal persistence.

Collectively, these findings establish metabolic dysregulation as a key driver of *C. auris* virulence and host susceptibility, providing a mechanistic framework linking nutrient sensing, immune dysfunction, and pathogen adaptation (**Figure 8**). This work highlights therapeutic opportunities aimed at targeting fungal adhesins and restoring host immune competence to prevent or limit *C. auris* colonization, particularly in diabetic and DKA patient populations, with important implications for surveillance, prevention, and antifungal intervention strategies.

**Figure 8:**
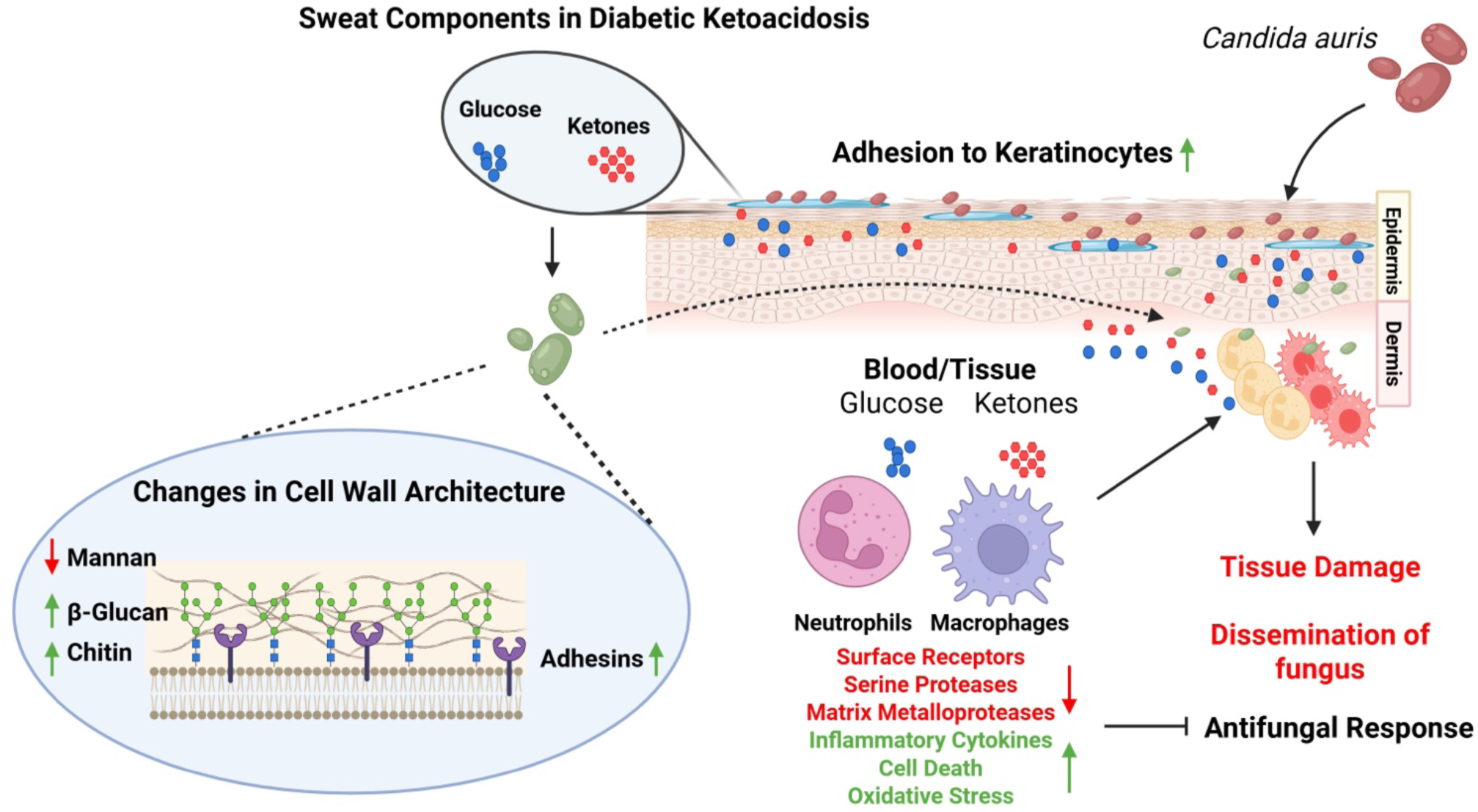
Proposed mechanistic insight of *C. auris* adhesion and increased virulence/colonization under diabetic ketoacidosis conditions.

## MATERIALS AND METHODS

### Isolates, Growth conditions, and Inoculum preparation

*C. auris* strains were acquired from Centers for Disease Control and Prevention. Chiacago4 strain provided by Dr. Teresa O’Meara at the University of Michigan^34^. Yeast strains were cultured overnight in Yeast Extract Peptone Dextrose (YPD) broth at 30 °C with continuous agitation at 200 rpm. Following incubation, cells were harvested and washed three times with 1× phosphate-buffered saline (PBS; Cat. No. 10010023, ThermoFisher Scientific). Blastoconidia were enumerated using a hemocytometer and adjusted to the desired cell concentration in specified media for subsequent experiments^33^. Preliminary strain screen demonstrated high adhesion capacity with the CAU-02, CAU-03, CAU-05, CAU-07 and Chicago4. We used Chicago4 strain in the entire study other than the adhesion assay. For growth assay, an overnight culture of Chicago4 and CAU-09 was washed twice with PBS and counted to maintain a cell concentration of 1x10^6^ CFU/ml. 10 μl of the fungal suspension was then grown in RPMI media supplemented with MOPS and either glucose alone (4 or 8 mg/ml), 3-hydroxybutyric acid (BHB) (10 or 20 mM, Cat: 166898, Millipore Sigma) alone or a combination of both, 37^0^C for 20 h under static condition. Absorbance was measured at 600 nm after every 30 min. For assays involving growth of *C. auris* in the presence of glucose and BHB, an overnight culture of either or both strains of *C. auris* was washed twice with PBS and a small fraction of the inoculum was sub-cultured in RPMI media in the presence of Glucose or BHB at 37^0^C overnight. The following day, the fungal cells were washed twice with PBS, diluted and counted to maintain CFU of requirement.

### *In vitro* fungal adhesion assay

We compared several *C. auris* isolates for their adhesion capability *in vitro* using XTT assay as previously described (**Supplementary Table S1**)^33,34,37,80^. Briefly, overnight cultures of yeast strains were washed twice with PBS and diluted to an inoculum size of 1x10^4^/ml in RPMI-1640 media containing MOPS (165 mM). 100 μl of yeast suspension was then added on to a 96 well plate and incubated overnight at 37^0^C. The following day, adhered fungal biofilm was gently washed twice with PBS, following which 100 μl XTT (2,3-Bis-(2-Methoxy-4-Nitro-5-Sulfophenyl)-2H-Tetrazolium-5-Carboxanilide) solution (Cat: X6493, ThermoFisher Scientific; 1 mg/ml) (containing 1 mM Menadione) was added to the adhered fungus. Post incubation at 37^0^C for 30-60 min and color development, the absorbance was read at 450 nm.

For adhesion to N/TERT-1 keratinocytes^61^, approximately 2 x 10^6^ keratinocytes were plated on a 6 well plate overnight in K-Sfm media supplemented with human recombinant epidermal growth factor and bovine pituitary extract (Cat: 17005042, ThermoFisher Scientific), 1% Pen-Strep antibiotic cocktail (Cat: 15140122, ThermoFisher Scientific) and 0.3 mM CaCl_2_ (Cat: AAJ63122AE, ThermoFisher Scientific). An overnight culture of Chicago4 grown under different conditions as described above was washed twice with PBS and maintained at concentration of 5 x 10^3^ CFU/ml in K-Sfm media. Prior to experiment, the media was removed from the N/TERT-1 keratinocytes^61^ and 200 μl of fungal inoculum was carefully laid over the cells. The fungus was allowed to adhere to the keratinocytes for 4 h at 37^0^C. The cells were gently washed twice with PBS to remove any planktonic fungus. YPD broth containing 0.8% agar was carefully poured on top of the keratinocytes and the plates were incubated at 37^0^C overnight. Similar volume of fungal inoculum was also plated on a YPD agar plate to serve as inoculum control. The following day CFU was counted on both control and adherence plate, and percentage adhesion was calculated thereafter.

### Establishment of a murine skin colonization model of *C. auris* infection

We used 5 to 6-week-old inbred C57BL/6 male mice weighing 25-30 grams purchased from Envigo, USA. Mice were kept either immunocompetent, made immunosuppressed or diabetic ketoacidosis (DKA) (n=5-6/group). Immunosuppression was induced by administering 200 mg/kg cyclophosphamide intraperitoneally (IP) and 20 mg/kg triamcinolone subcutaneously (SC) 2 days before the start of the *C. auris* inoculation. The immunosuppression was maintained throughout the experiment by administering cyclophosphamide (100 mg/kg) and triamcinolone (10 mg/kg) every 5 days. Immunosuppression was confirmed by counting total WBCs in the blood using Leuko-TIC WBC counting kit (Bioanalytic GmbH).

Diabetic Ketoacidosis (DKA) was induced by a single IP injection of 175 mg/kg streptozotocin (SZT) (Cat. No. 572201, Millipore Sigma), dissolved in phosphate citrate buffer, at 10 days before *C. auris* inoculation^81^. High blood glucose levels (>300 mg/dL) in mice were confirmed using automated glucose testing strips, whilst ketonuria was confirmed using Ketostix Urine Test Strips on day 7 post SZT treatment. To prevent bacterial superinfection in immunosuppressed and DKA mice, enrofloxacin (Baytril®, Elanco, Germany) was administered via drinking water at a concentration of 50 µg/mL, starting on the day of infection and continuing through day 10 post-infection.

The shaved backs of anesthetized mice (2 cm^2^ area) were repeatedly inoculated for 4 days with 0.1 ml artificial sweat media containing 10^9^ CFU of *C. auris* blastoconidia, spread with sterile cotton swabs, and air-dried for 2 minutes. Colonization was monitored by sampling of inoculated skin surface on days 2 (baseline), 7, 14, and 21 using sterile cotton swabs. The colonized skin was imaged and compared among immunocompetent, immunosuppressed, and DKA mice. On days 2, 7, and 21, skin tissues and kidney tissues (day 21 only) for systemic dissemination were assessed following euthanization (**Figure 1a**).

### Determination of skin fungal burden

For fungal burden determination, cotton swabs with 1 ml diluent (1X PBS), weighed skin or harvested kidney tissue were homogenized and quantitatively cultured using 10-fold serial dilutions on CHROMagar^TM^ Candida Plus (Cat: CA242, CHROMagar) plates supplemented with chloramphenicol. The plates were incubated at 37°C for 48 hours before colony-forming units (CFUs) per gram of tissue or per ml (for swabs only) were enumerated ^33,35^.

### Skin Histopathology

For the histopathological examination, skin biopsies from representative mice were harvested on days 3 and 7 post-infection. The skin was fixed in 10% zinc-buffered formalin, dehydrated using ethanol and xylene, embedded in paraffin, sectioned, and stained with PAS staining kit (Cat: 1.01646, Millipore Sigma) and Hematoxylin. Stained sections were examined and imaged using an Olympus microscope.

### Primary Mammalian Cell Culture

Bone marrow was isolated from femurs and tibia of uninfected Immunocompetent and DKA mice. Bone marrow neutrophils (BMN) were purified using the MojoSort Mouse Neutrophil Isolation Kit (Cat: 480058, BioLegend) according to the manufacturer’s instructions. Bone marrow cells were differentiated into primary macrophages *in vitro* for 6 days using GM-CSF (20 ng/ml) in RPMI-1640 media supplemented with L-Glutamine (Cat: MT10041CM, Fisher Scientific) and 10% FBS (Cat: A5670801, ThermoFisher Scientific), 1% Pen-Strep antibiotic cocktail (Cat: 15140122, ThermoFisher Scientific).

### Phenotyping of immune cells by flow cytometry

Immune cell infiltration was evaluated in skin tissue homogenates by flow cytometry. Skin biopsies were collected on days 3 and 7 post-infection. The tissues were homogenized by enzymatic treatment using a cocktail of Liberase^TM^ (TL Research Grade, Cat: 5401020001, Milipore Sigma) and DNAse I (Cat: DN25, Milipore Sigma) at 0.25 mg/ml and 1, μg/ml respectively, in RPMI (not supplemented with antibiotics or FBS)^83^ for 2 h at 37^0^C. Following enzymatic digestion, the reaction was stopped using complete RPMI. The skins were crushed using the plunger of a 1 mL syringe, and a single-cell suspension was prepared by passing the skin homogenate through a 70 μm filter. The tissue homogenate containing immune cells was then stained with fluorescent antibodies against CD3, CD4, CD19, F4/80, CD11b, CD11c, and Ly6G. The details of the antibodies used are provided in **Supplementary Table S2**.

Neutrophil and resident macrophage populations in the bone marrow of immunocompetent and DKA mice were quantified by flow cytometry. Briefly, bone marrow cells, resuspended in Lift Buffer (PBS containing 2% FBS, 0.5 mM EDTA and 0.1% Sodium azide) were stained for Ly6G, CD11b and F4/80 for an hour at 4^0^C. All stained cells were acquired using a BD LSR II flow cytometer, and the data were analyzed with FlowJo V10.

### Determination of fungal cell wall components by flow cytometry

Cell wall components of *C. auris* were determined by flow cytometry as previously described ^56^. Briefly, an overnight culture of Chicago4 and CAU-09 was grown in the presence or absence of glucose or BHB or both as described above. Fungal inoculum was washed twice with PBS and fixed with 10% Zinc Formalin at 37^0^C for 1 h. The inoculum was washed twice with PBS and then stained either for mannan with Concanavalin A (50 μg/ml, Cat: C860, ThermoFisher Scientific) for 1 h or for chitin with Calcofluor White (30 μg/ml, Cat: 18909, Millipore Sigma) for 10 min at 37^0^C. Fungal cells were washed twice with PBS before analyzing cell wall components using flow cytometry as described above.

### Determination of cytokines from skin biopsy and cell culture supernatants

Skin tissues collected on days 3 and 10 post-inoculation were homogenized as described above. Single cell suspension of skin cells was lysed using RIPA buffer supplemented with 1 mM PMSF and 1X Protease Inhibitor Cocktail (Cat: PI78429, ThermoFisher Scientific). Cell debris was removed by centrifuging the lysate at 14000 g at 4^0^C for 5 min. Cell lysate supernatant was used to determine inflammatory cytokines using the LegendPlex^TM^ Mouse Inflammation Panel (13-Plex) Kit (Cat: 740446, BioLegend) and flow cytometry. Data was analyzed using the LegendPlex Software. Bone Marrow Derived Macrophages (BMDM) from immunocompetent and DKA mice were infected with *C. auris* (Chicago4, MOI: 5) for 4 h. The supernatant was collected and centrifuged at 4000 g for 5 min to remove any fungus. The supernatant was then analyzed using the LegendPlex kit for release of pro-inflammatory cytokines. For determination of specific cytokines, BMNs and BMDMs were infected with Chicago4 (MOI:5), grown under different conditions as described above, for 4 h. Cell culture supernatants were centrifuged to remove any planktonic fungus, and then the supernatant was analyzed for secreted levels of TNF- α (Cat: DY410, R&D Systems) and IL-6 (Cat: DY406, R&D Systems). To detect IL-1β, infected cells were further treated for 1 h with 10 μM Nigericin (Cat: N7143, Millipore Sigma), following which, the supernatant was harvested and analyzed for the cytokine (Cat: DY401, R&D Systems).

### Determination of Phagocytic uptake of labelled *C. auris*

Uptake of *C. auris* by phagocytes was determined by flow cytometry as previously described ^85^. Briefly, an overnight culture of Chicago4 strain (grown with or without glucose and BHB) was washed twice with PBS and then manually counted on a hemocytometer to adjust the MOI to 1. Fungal cells were then stained with 10 μM CellTrace CFSE Cell Proliferation Kit (Cat: C34554, ThermoFisher Scientific) for 45 min at 37^0^C. The stained fungal cells were washed twice with PBS to remove any additional dye. Approximately, 1x10^6^ phagocytes (BMNs and BMDMs) were then infected with the labelled fungus for 1.5 h. Cells were then washed twice with PBS to remove any planktonic *C. auris*. The cells were resuspended in Lift Buffer, and fungal uptake was assessed using flow cytometry.

### Determination of total cellular Reactive Oxygen Species (ROS)

Approximately, 1x10^6^ phagocytes (BMNs and BMDMs) were pre-stained with 10 μM CM-H_2_DCFDA (Cat: C6827, ThermoFisher Scientific) for 30 min at 37^0^C. The cells were washed twice with PBS to remove any excess dye and then infected with an overnight culture of Chicago4, maintained at MOI-1, for 1 h. The cells were washed twice with PBS to remove any planktonic fungi and then resuspended in Lift Buffer. Total cellular ROS was then assessed using flow cytometry^85,86^.

### Flow Cytometric analysis of the expression of cell surface receptors

Approximately 1x10^6^ unstimulated BMDMs from Immunocompetent and DKA mice were stained with fluorescently labelled antibodies against cell surface receptors including Dectin-1 (CLEC7A), TLR2 and TLR4 for 1 h at 37^0^C. Post staining, the cells were washed twice with PBS to remove any excess unbound antibodies. Finally, the cells were resuspended in Lift Buffer and cell surface receptor expression was assessed using flow cytometry. Details of the antibodies used are listed on **Supplementary Table 2**.

### Determination of phagocyte killing exposed to *C. auris*

Cytotoxicity of BMNs and BMDMs (1x10^5^ cells/well) post infection with Chicago4 (MOI-5, 4 h) was determined by measuring lactate dehydrogenase in the cell culture supernatant using the CytoTOX 96 Non-Radioactive Cytotoxicity Assay Kit (Cat: G1780, Promega), as per the manufacturer’s instructions.

### Determination of fungal killing by phagocytes

Approximately 1x10^6^ BMNs and BMDMs were seeded onto a 24-well plate in complete RPMI-1640 media. The cells were allowed to settle for 2 h before infecting them with an overnight culture of Chicago4, grown in the presence or absence of glucose and BHB (MOI-5, 4h). To determine percentage killing, Chicago4 was also cultured in the absence of mammalian cells in complete RPMI for 4 h. Host cells were then lysed using 0.05% Triton X-100. Number of viable fungal cells in the presence or absence of host cells was determined by plating the fungus on YPD agar plates and counting the CFU the following day as previously described^87^.

### Determination of mammalian and fungal gene expression

Approximately 4x10^6^ BMNs or 1x10^6^ BMDMs from Immunocompetent and DKA mice were either left uninfected or were infected with an overnight culture of Chicago4 (MOI-5) for 4 h. The host cells were washed once with PBS to remove any planktonic fungus and then total RNA was extracted using RNA purification kit (Cat: 74534, Qiagen), as per the manufacturer’s instructions. DNA contamination was removed by treating the RNA samples with DNAse I (Cat: EN0521, ThermoFisher Scientific) as per the manufacturer’s instructions. RNA was reverse transcribed into cDNA using SuperScript^TM^ IV VILO^TM^ Master Mix (Cat: 11756050, ThermoFisher Scientific). Levels of specific mRNA were quantified by qPCR using PowerTrack^TM^ SyBR Green Master Mix (Cat: A46109, ThermoFisher Scientific) in the Applied Biosystems StepOne Plus Real-Time qPCR system. Appropriate negative controls with no SuperScript were included for all experiments. Data were expressed relative to the housekeeping gene, hypoxanthine phosphoribosyltransferase (*Hprt*, mouse) using the ΔΔCt method and fold change was calculated relative to uninfected BMN or BMDM from Immunocompetent mice. To determine expression of adhesin gene from fungus, total RNA was isolated from Chicago4 using Direct-zol RNA Kit (Cat: R2053, Zymo Research), as per the manufacturer’s instructions. Briefly, an overnight culture of Chicago4 was grown in the presence or absence of glucose and BHB as described previously. Fungal cells were washed twice with PBS and then resuspended in TRI Reagent and frozen at -80^0^C overnight. The following day, fungal cells were lysed by bead-beating method using Lysing Matrix Y (Cat: SKU:116960100, MP Biomedicals) and the total RNA was extracted. DNA contamination was removed and total RNA was reverse transcribed into cDNA as described above. Levels of specific mRNA were then quantified using qPCR. Appropriate negative controls with no SuperScript were included for all experiments. Data were expressed relative to the housekeeping gene, β-actin (*Act*, *C. auris*) using the ΔΔCt method and fold change was calculated relative to control Chicago4. A list of all the primers used in this study is provided on **Supplementary Table S3**.

### Serum Antibody Binding to Fungal Cell Surface

An overnight culture of Chicago4 and CAU-09 were grown in the presence or absence of glucose and BHB as described above. Fungal cells were washed twice with PBS and fixed as described above. The fungal inoculum was then washed twice with PBS and blocked with 3% bovine serum albumin (Cat: 10735078001, Millipore Sigma) for 1 h at 37^0^C. The cells were pelleted and resuspended in PBS containing sera (1:500) from either control or NDV-3A vaccine (containing *Candida albicans* N-terminal recombinant Als3p protein)^33^ or (*C. albicans* recombinant N-terminal protein) Hyr1 protein vaccinated mice ^35,64^. The cells were incubated for 2 h at 37^0^C, washed twice with PBS and then resuspended in PBS containing Alexa Fluor 488 labelled anti-mouse IgG detection antibody (Cat: A-24920, ThermoFisher Scientific). Post 1 h incubation, the cells were washed twice with PBS before analyzing them by flow cytometry as described above.

### Determination of Neutrophil Elastase

Approximately 2x10^6^ BMNs from Immunocompetent and DKA mice were either left uninfected or were infected with Chicago4 (MOI-5) for 4 h. The supernatant was collected and centrifuged to remove any planktonic fungus. BMNs were then lysed using RIPA buffer containing PMSF and Protease Inhibitor Cocktail. Neutrophil elastase in the supernatant as well as the lysate were then determined using the NETosis Assay Kit (Cat: 601010, Cayman Chemical) as per the manufacturer’s instructions.

### Statistical analysis

Sample size for fungal burden experiments was calculated assuming an α of 0.05, 90% power, and a minimum detectable effect size of 1.0 log CFU, yielding an estimated requirement of approximately five mice per group.

All *in vitro* experiments were performed at least twice with multiple technical replicates. *In vivo* and *ex vivo* experiments were conducted with at least 5 mice per group. Data for fungal burden, immune cell counts, and cytokine levels are presented as median ± interquartile range (IQR). Differences between the two treatment groups were evaluated using the Mann–Whitney test, while multiple group comparisons for cytokine analysis employed the Kruskal–Wallis test followed by Dunnett’s post hoc correction. These nonparametric tests were selected to minimize the influence of outliers and are appropriate for small sample sizes.

To generate heatmaps for gene expression, fold changes were first transformed into their logarithmic values. Difference in fold change were assessed using Kruskal–Wallis test or two-way ANOVA followed by Dunn’s and Dunnett’s post hoc correction respectively.

Graphical representation and statistical analyses were performed using GraphPad Prism (v10.4). Detailed descriptive statistics are provided in the main figures or supplementary materials. The histopathological images shown are representative of more than three biological replicates.

## ACKNOWLEDGMENTS

We acknowledge Dr. Teresa O’Meara (University of Michigan, Ann Arbor, MI, USA) for providing *C. auris* Chicago4 strain. We also thank The Brigham and Women’s Hospital, Inc for providing N/TERT-1 cell lines.

## DATA AND MATERIALS AVAILABILITY

All data from this study are available in the main text or the supplementary materials.

## AUTHOR CONTRIBUTIONS

Conceptualization: S.S.; Funding acquisition: S.S.; Investigation: K.D.G.; Methodology: K.D.G, D.Q., E.G.Y., K.G., S.S.; Project administration: S.S.; Supervision: S.S.; Data collection: K.D.G., Visualization: K.D.G, S.S.; Writing – original draft: S.S., K.D.G.; Critical inputs and Material: A.S.I., Writing – review and editing: K.D.G., A.S.I., S.S.

