## Supplementary material for "Diabetes and Immunosuppression Drive Distinct Patterns of *Candidozyma auris* Skin Colonization and Dissemination in Mice": Supplimentary Material

*Corresponding author

**Short Title**: Host metabolic and immune status drives *Candidozyma auris* colonization

**Short Summary**: This study establishes the first physiologically relevant murine model of *Candidozyma auris* skin colonization under diabetic ketoacidosis and immunosuppression, revealing distinct immune dysfunction and systemic dissemination that can inform targeted antifungal strategies

**a.**

| Day | Blood Glucose ± SD mg/dl in Mice | | |
|---|---|---|---|
| | Immuno-competent | Immuno-suppressed | Diabetic-ketacidosis |
| -6 | 226.7 ± 30.99 | 213.2 ± 32.06 | >500 (13/17) |
| -3 | 199 ± 32.06 | 196.4 ± 26.87 | >500 (17/17) |
| | Urine Ketone levels (mg/dl) in Mice | | |
| -6 | Undetectable | Undetectable | >160 mg/dl (17/17) |
| -3 | Undetectable | Undetectable | >160 mg/dl (17/17) |

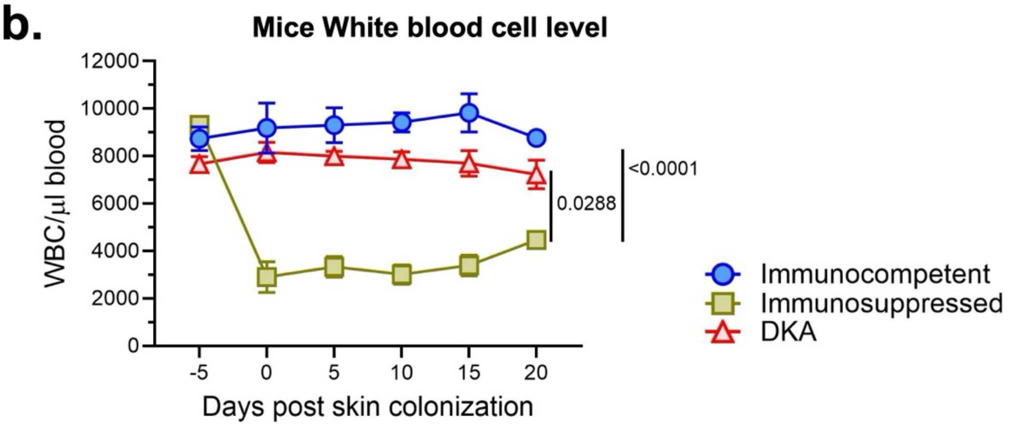

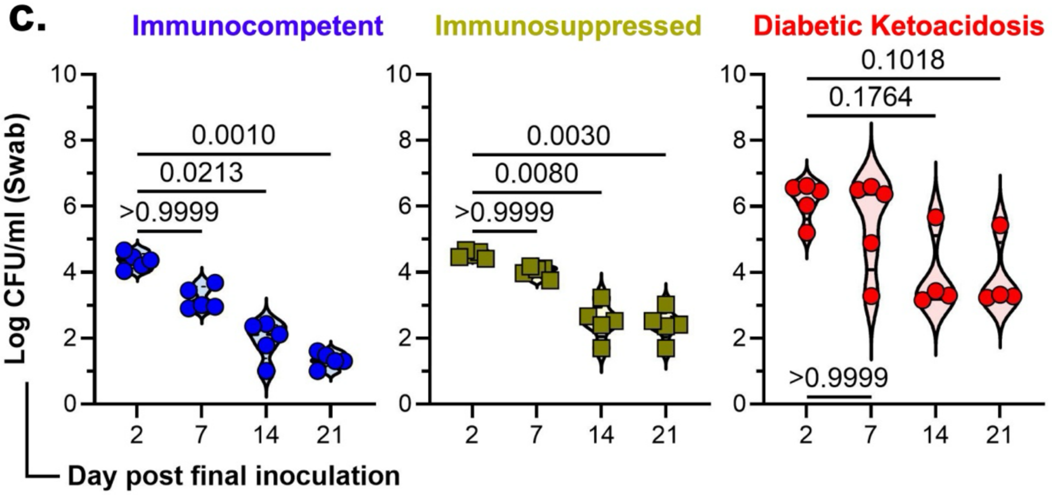

**Figure S1 (Related to Figure 1). : Validation of Immunosuppression and Diabetes induction in mice and change in skin swab fungal load with time.** (**a**) *Ad libitum* blood glucose and urine ketone level estimation in mice from different groups. (**b**) WBC count in mice from different groups across the period of skin colonization. Statistical analysis was performed using a two-way ANOVA followed by Tukey's multiple-comparison test. (**c**) *C. auris* skin colonization monitoring over time compared across the different clinical groups. Data are represented as violin plots and expressed as median $\pm$ interquartile range. Statistical analysis was performed using the Kruskal-Wallis test followed by Dunn's multiple comparison test. For statistical significance, a p-value less than 0.05 was considered significant.

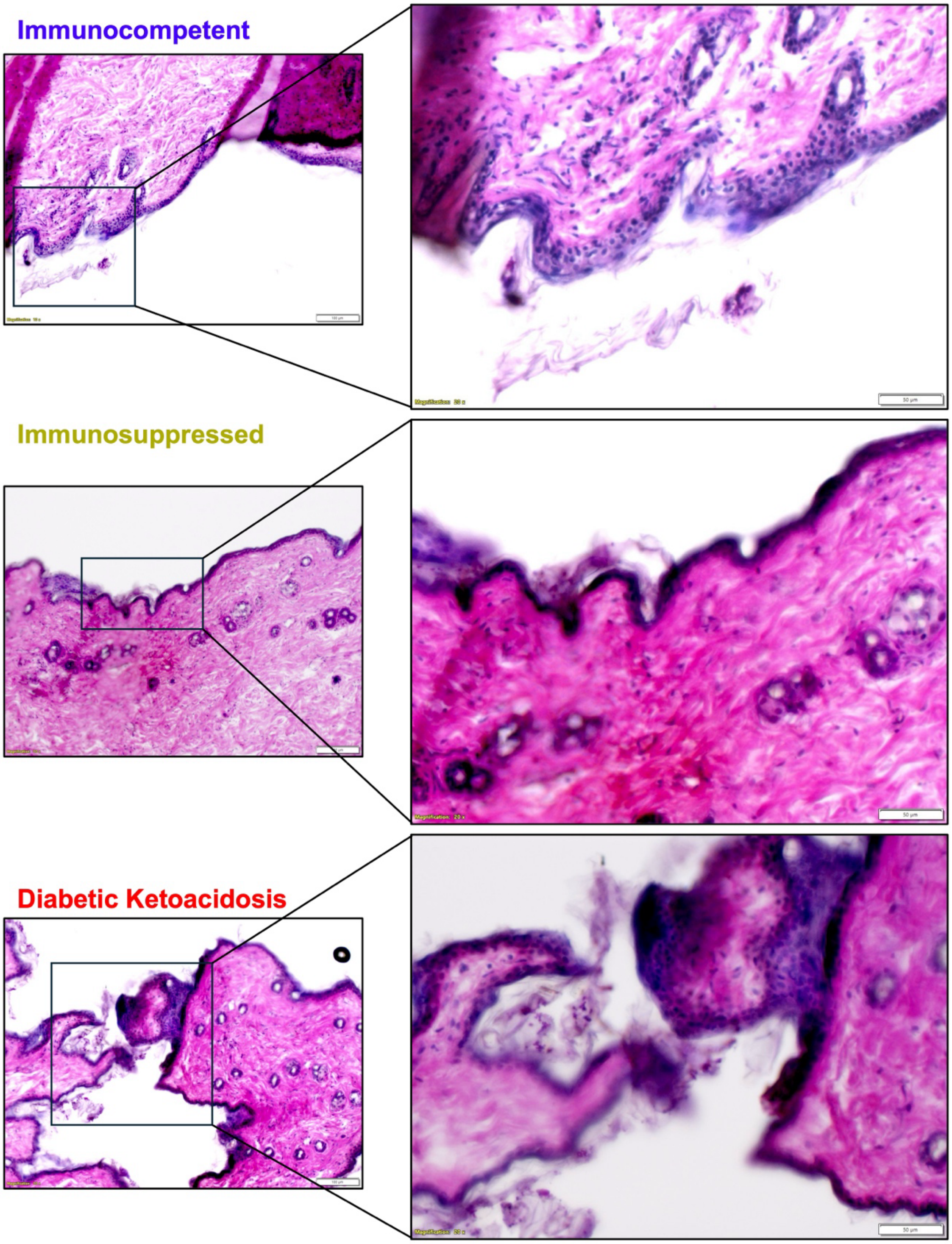

**Figure S2 (Related to Figure 2). Histopathology of the skin from immunocompetent, immunosuppressed and DKA mice**. Histopathology were stained and imaged on day 7 post *C. auris* skin colonization (Scale bar: 10 μm) with zoomed insets (Scale bar: 20 μm) are presented.

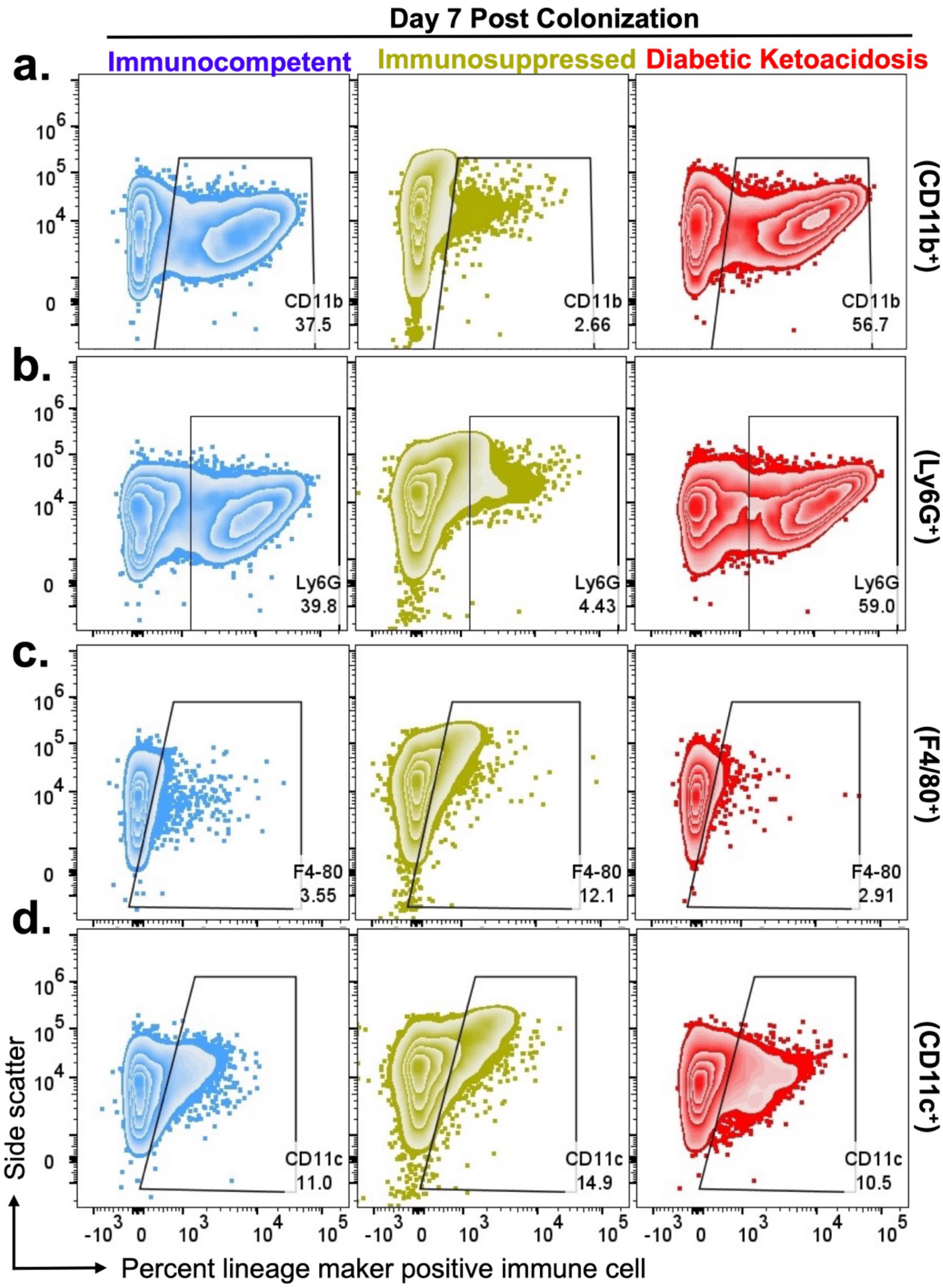

**Figure S3 (Related to Figure 3). Immunophenotyping of *C. auris* colonized skin tissues.** (**a-d**) Representative scatter graphs showing infiltrating immune cell population in the skin 7-days post colonization of immunocompetent, immunosuppressed and DKA mice. The stained cells were analyzed using BD FACSymphony, and the area was analyzed with FlowJo v10.

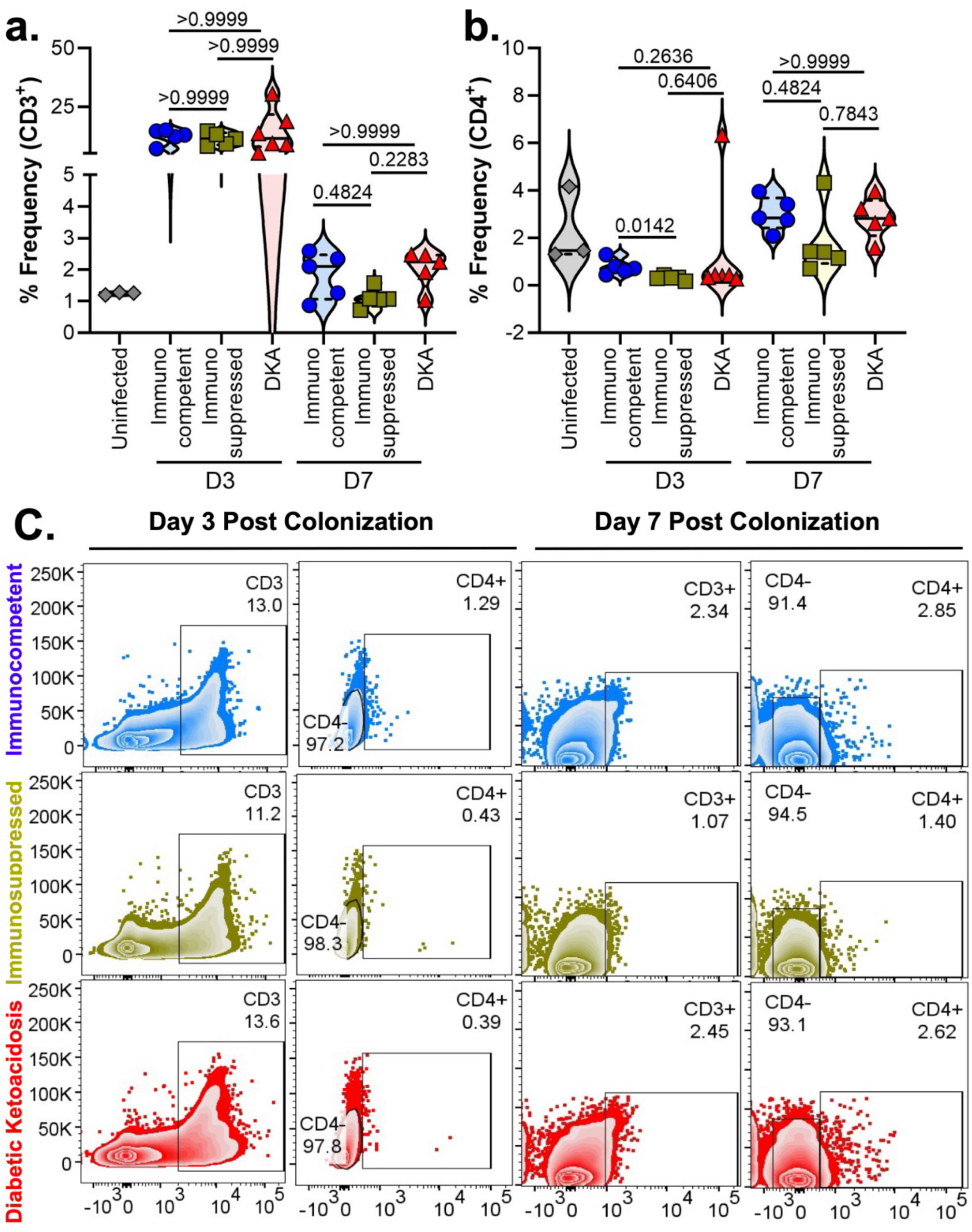

**Figure S4 (Related to Figure 3). T cell profiling of *C. auris* colonized skin tissues.** (**a-c**) Flow Cytometric determination of percentage CD3 and CD4 cells in the skin 3- and 7-days post colonization of immunocompetent, immunosuppressed and DKA mice. The stained cells were analyzed using BD FACSymphony, and the area was analyzed with FlowJo v10. Representative scatter graphs and corresponding violin plots (median ± interquartile range) are shown for the different clinical groups. Significant differences in the immune cell populations were analyzed by the Kruskal-Wallis test, followed by Dunn's multiple comparison test. p values <0.05 were considered statistically significant.

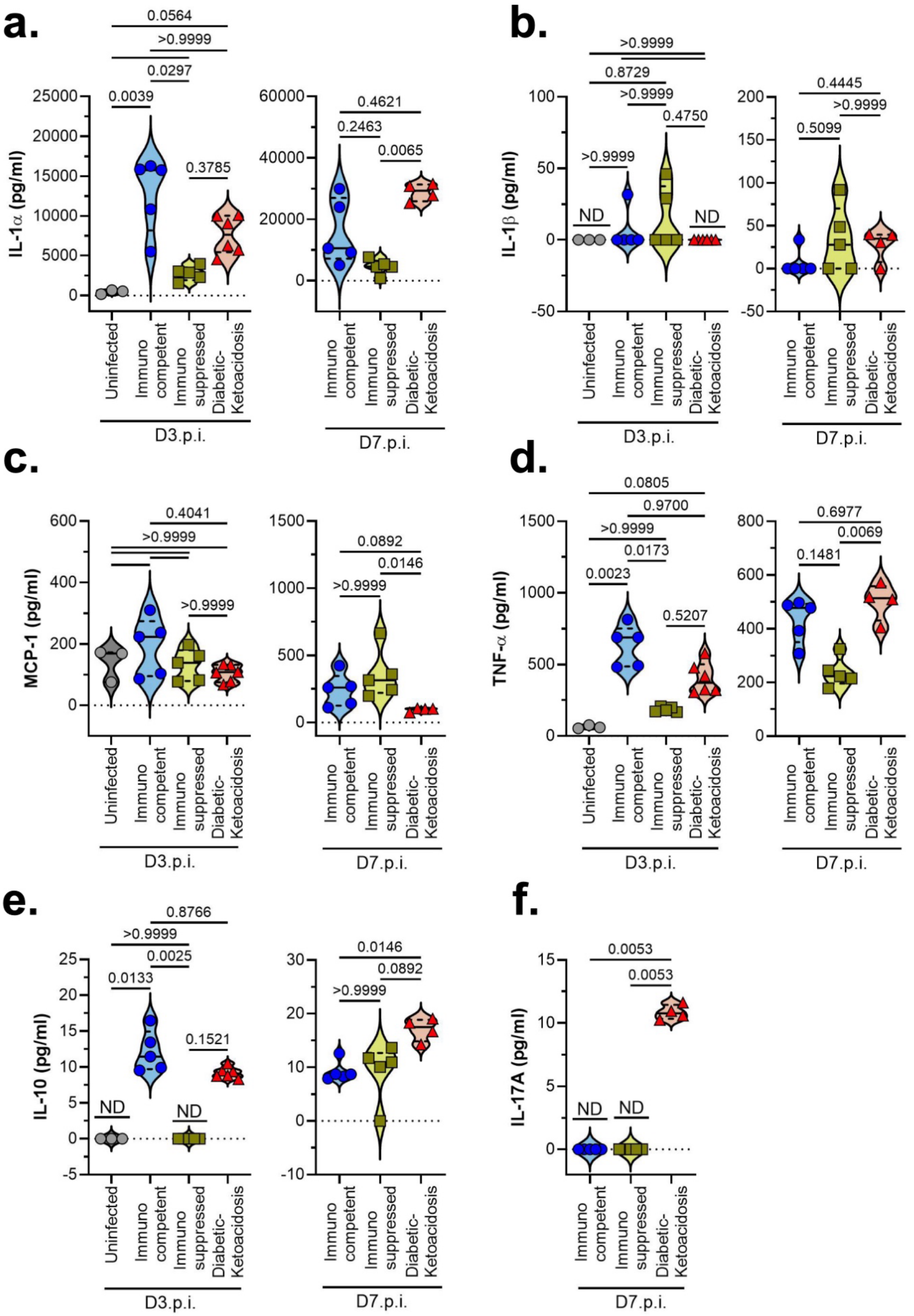
a.
IL-1α (pg/ml)
0.0564
>0.9999
0.0297
0.0039
0.3785
0.4621
0.2463
0.0065
Uninfected
Immuno competent
Immuno suppressed
Diabetic-Ketoacidosis
D3.p.i.
D7.p.i.
b.
IL-1β (pg/ml)
>0.9999
0.8729
>0.9999
0.4750
>0.9999
ND
ND
0.4445
>0.9999
0.5099
c.
MCP-1 (pg/ml)
0.4041
>0.9999
>0.9999
0.0892
0.0146
>0.9999
d.
TNF-α (pg/ml)
0.0805
0.9700
>0.9999
0.0173
0.0023
0.5207
0.6977
0.0069
0.1481
e.
IL-10 (pg/ml)
0.8766
>0.9999
0.0025
0.0133
0.1521
ND
ND
0.0146
0.0892
>0.9999
f.
IL-17A (pg/ml)
0.0053
0.0053
ND
ND

**Figure S5. Differential inflammatory cytokine responses in *C. auris* colonized skin immunocompetent, immunosuppressed, and DKA mice.** Immunocompetent, Immunosuppressed and DKA mice (n=5/group) were colonized with 1 × $10^9$ yeast cells/mouse of *C. auris* (Chicago4) for 4 d. 3- and 7-days post colonization, the colonized skin was enzymatically digested and processed to prepare a single-cell suspension and lysed to release all intracellular cytokines. Cytokine concentration was determined by cytokine bead-array. (**a-f**) Intracellular levels of IL-1$\alpha$, IL-1$\beta$, MCP-1, TNF-$\alpha$, IL-10 and IL-17A are represented as violin plots. Data are expressed as median $\pm$ interquartile range. Statistical analysis was performed using the Kruskal-Wallis test followed by Dunn's multiple comparison test. For statistical significance, a p-value less than 0.05 was considered significant.

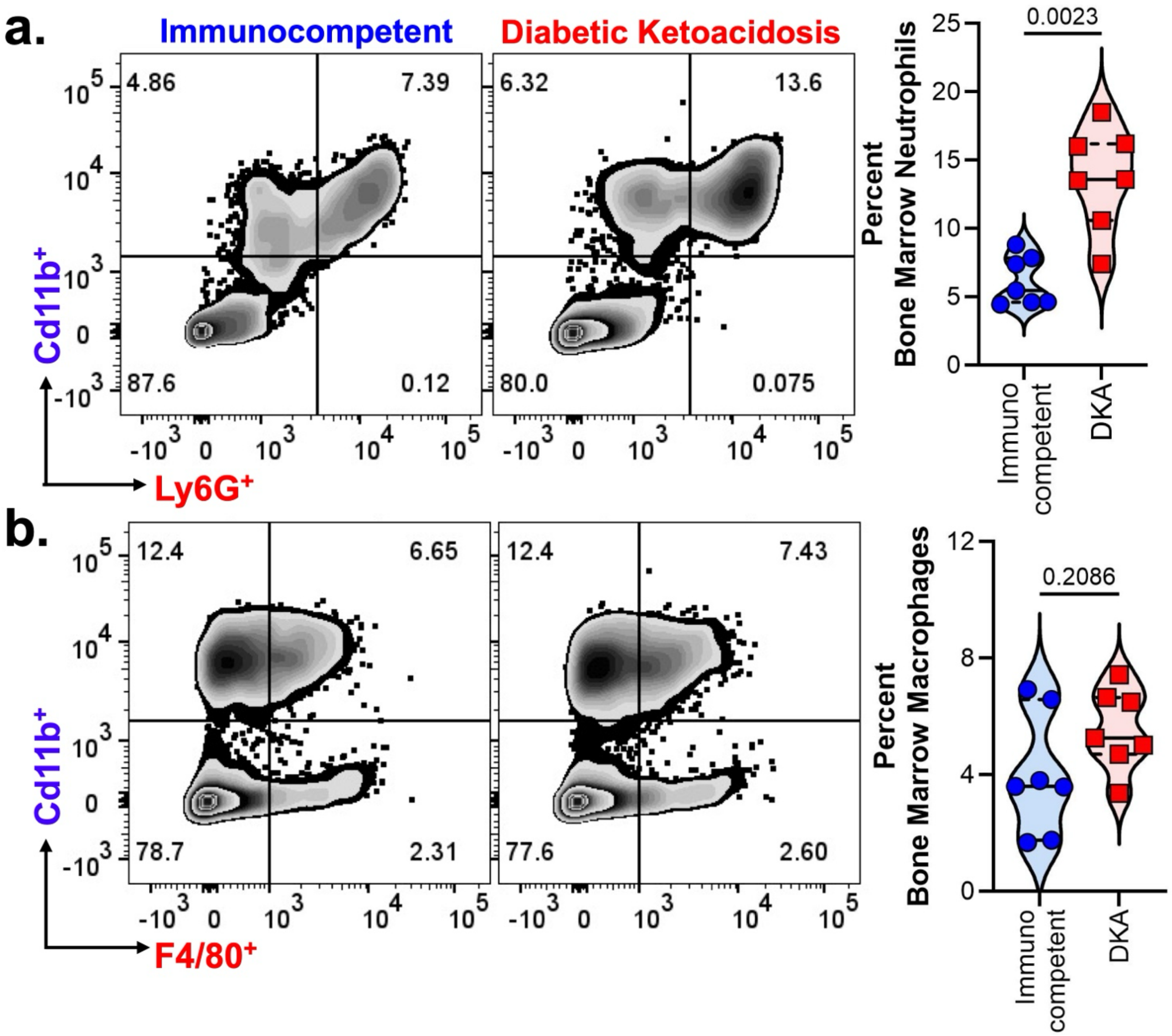

**Figure S6 (Related to Figure 4). Bone marrow profiling in Immunocompetent and DKA mice.** (**a-b**) Flow cytometric determination of percentage neutrophils (Ly6G$^+$ CD11b$^+$) and macrophages (F4/80$^+$ CD11b$^+$) in the bone marrow of immunocompetent and DKA mice. Representative scatter graphs and corresponding violin plots (median $\pm$ interquartile range) are shown for the different clinical groups. Significant differences in the immune cell populations were analyzed by the two-tailed Mann Whitney test. p values <0.05 were considered statistically significant.

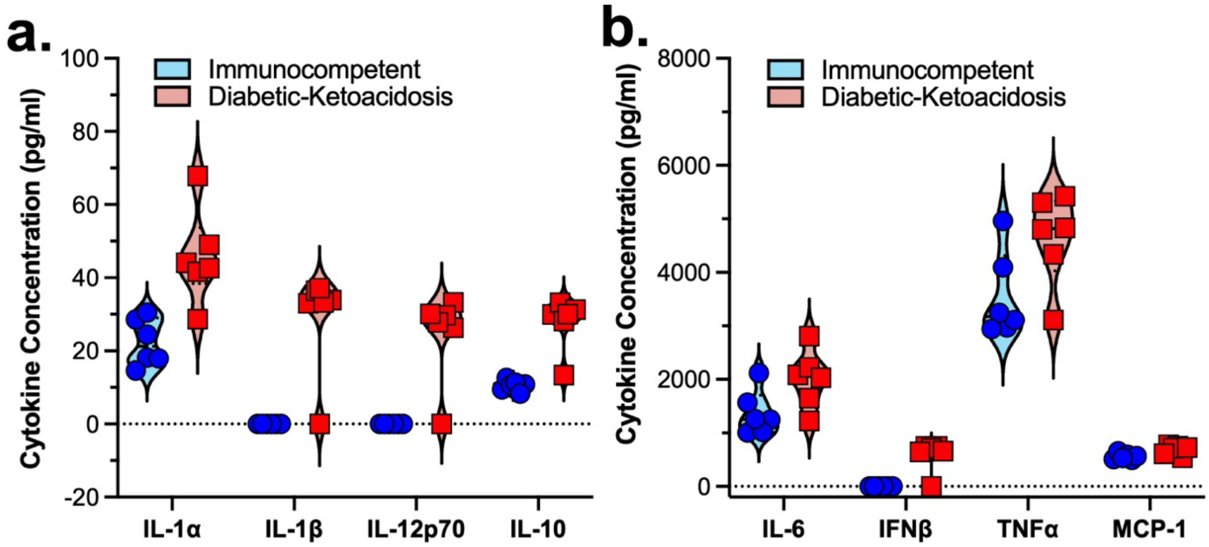

**Figure S7 (Related to Figure 5). Elucidation of cytokine release from Bone marrow derived macrophage from Immunocompetent and DKA mice upon infection with *C. auris*.** (**a-b**) Violin plots showing the cytokine concentration in the cell culture supernatant of primary macrophages from immunocompetent and DKA mice. Data are represented as median $\pm$ interquartile range.

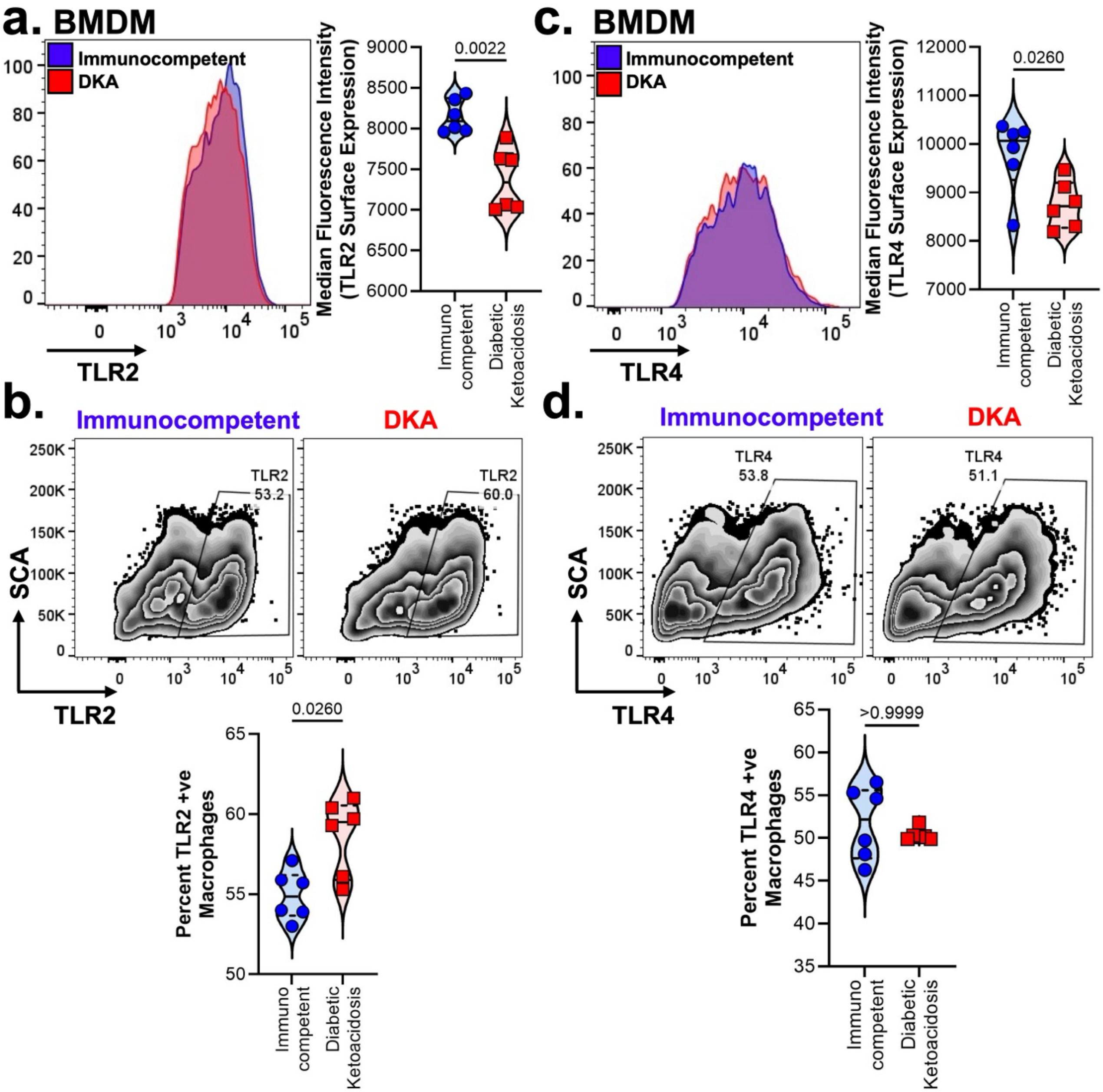

**Figure S8 (Related to Figure 6). Surface expression of TLR2 and TLR4 in Bone marrow derived macrophage from Immunocompetent and DKA mice .** (**a-d**) Surface expression of TLR2 (**a-b**) and TLR4 (**c-d**) in primary macrophages was determined by flow cytometry. Representative histogram plot and calculated median fluorescence intensity are presented (**a, c**). Representative zebra plot and calculated percentage *C. auris* positive neutrophils are presented (**b, d**). All violin plots are expressed as median $\pm$ interquartile range. Statistical analysis was performed using the two-tailed Mann Whitney test. For statistical significance, a p-value less than 0.05 was considered significant.

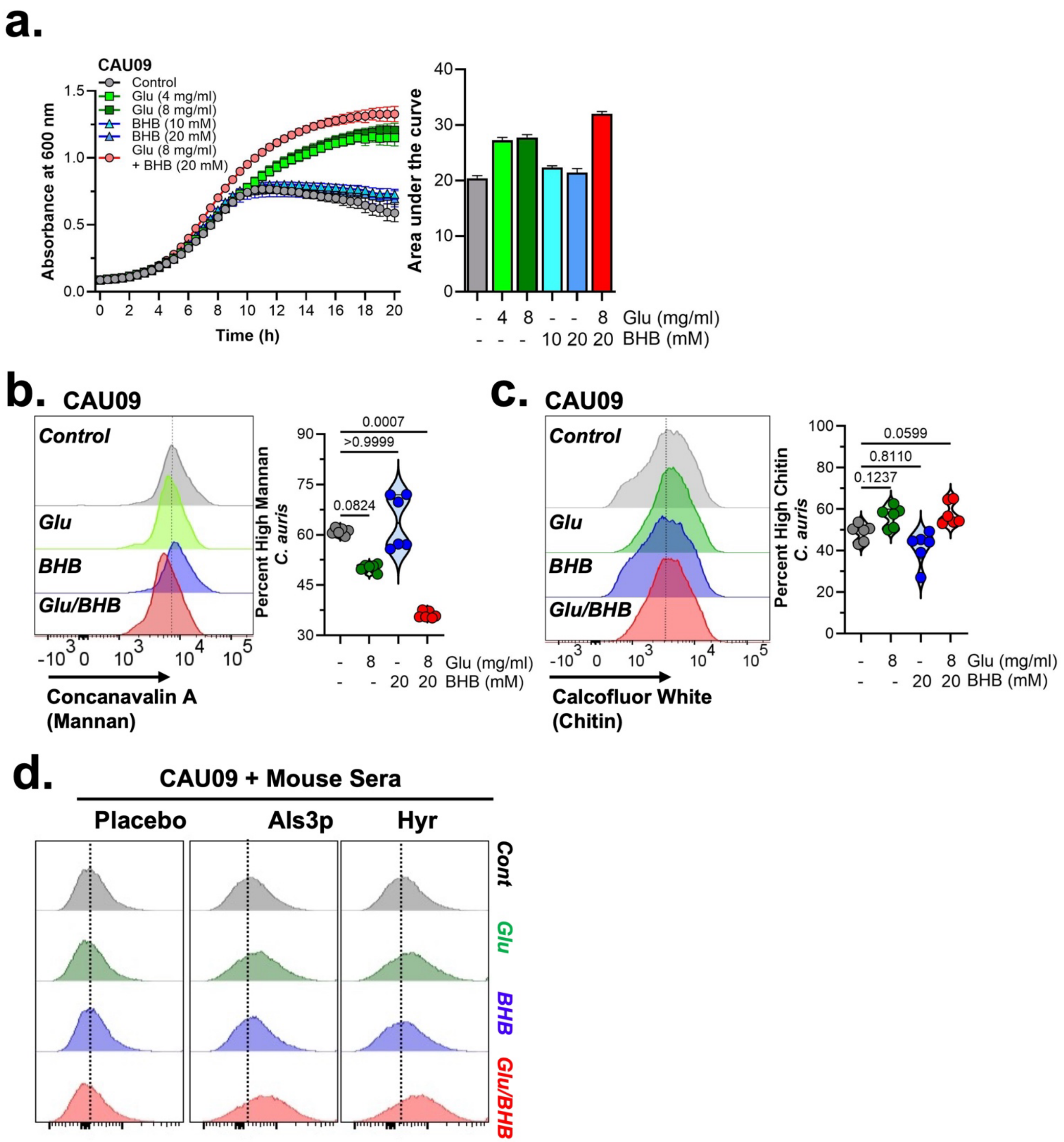

**Figure S9 (Related to Figure 7). Hyperglycemia and Ketoacidosis modulate *C. auris* cell wall architecture.** (**a**) The clade I strain CAU-09 was grown in the presence or absence of glucose or BHB alone or in combination for 20 h. Absorbance was measured at 600 nm as a measure of growth. Area under the curve was calculated thereafter. (**b-d)** CAU-09 was grown in the presence or absence of glucose or BHB alone or in combination overnight. Fungal cells were fixed and cell wall was stained for mannan (**b**) and chitin (**c**). Expression of these cell wall components were then analyzed by flow cytometry. Representative histogram plot and calculated percentage high cell wall components are presented. (**e**) Fixed CAU-09 cells grown under the high glucose and BHB conditions were incubated with control, anti-Als3p or anti-Hyr1p mouse sera. Antibody binding to fungal cell wall was detected with Mouse IgG antibodies conjugated with Alexa Fluor 488 using flow cytometry. Representative histogram plots are presented. All violin plots are expressed as median $\pm$ interquartile range. Statistical analysis for (**b-c**) was performed using Kruskal-Wallis test followed by Dunn's multiple comparison test. For statistical significance, a p-value less than 0.05 was considered significant.

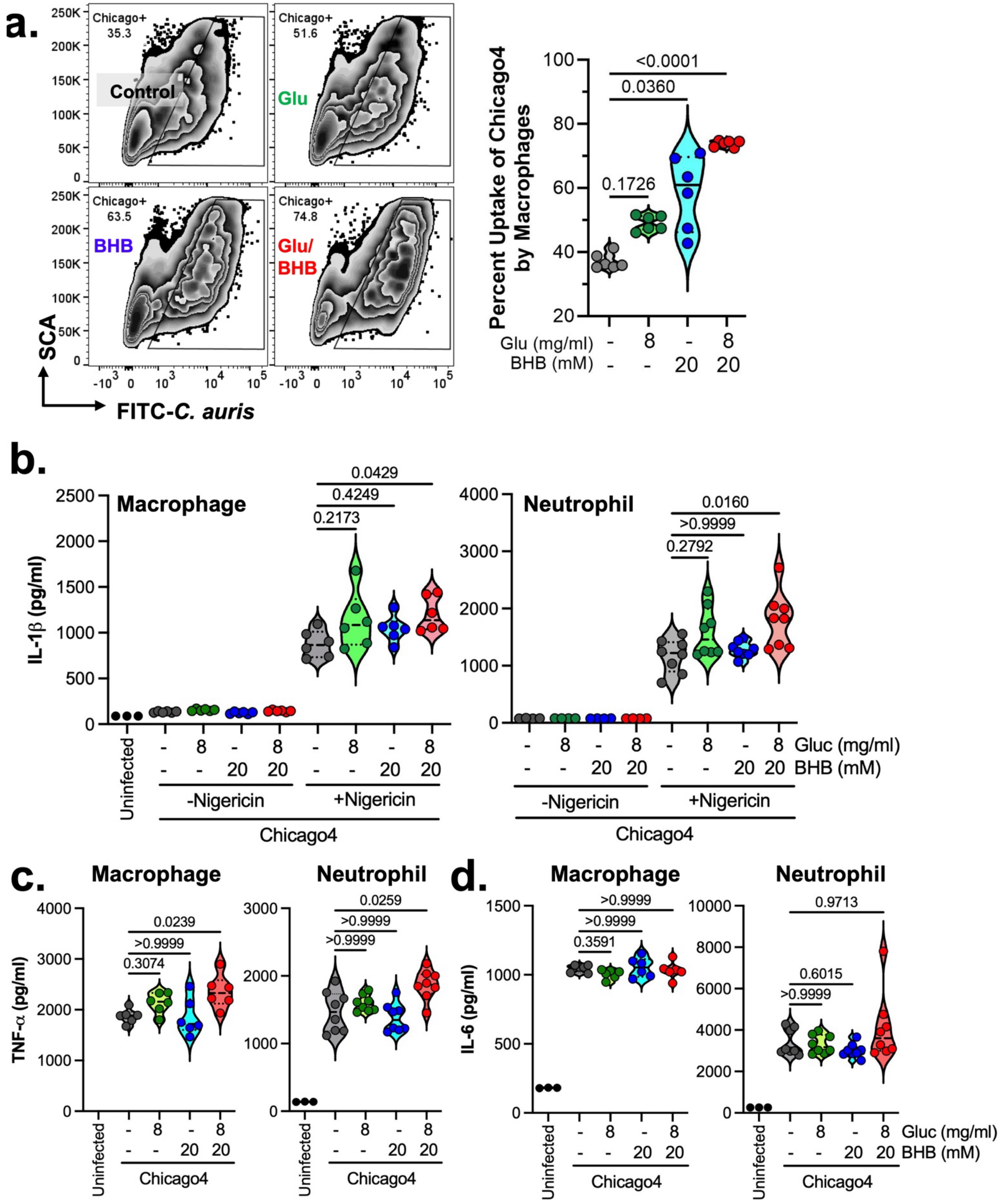
a.
Chicago+ 35.3
Control
Chicago+ 51.6
Glu
Chicago+ 63.5
BHB
Chicago+ 74.8
Glu/ BHB
SCA
FITC-C. auris
Percent Uptake of Chicago4 by Macrophages
<0.0001
0.0360
0.1726
Glu (mg/ml) - 8 - 8
BHB (mM) - - 20 20
b.
Macrophage
IL-1β (pg/ml)
0.0429
0.4249
0.2173
Neutrophil
0.0160
>0.9999
0.2792
Uninfected
Gluc (mg/ml)
BHB (mM)
-Nigericin
+Nigericin
Chicago4
c.
Macrophage
TNF-α (pg/ml)
0.0239
>0.9999
0.3074
Neutrophil
0.0259
>0.9999
>0.9999
d.
Macrophage
IL-6 (pg/ml)
>0.9999
>0.9999
0.3591
Neutrophil
0.9713
0.6015
>0.9999
Uninfected
Chicago4
Gluc (mg/ml)
BHB (mM)

**Figure S10 (Related to Figure 7). Hyperglycemia and Ketone rich condition modulate *C. auris* cell wall architecture dictate downstream inflammatory and antifungal responses of innate immune cells.** (**a**) Uptake of Chicago4, under different conditions, by BMDMs was determined by flow cytometry. Representative zebra plots and calculated percentage uptake are presented. (**b-d**) IL-1$\beta$, IL-6 and TNF-$\alpha$ released in the culture supernatant of neutrophils and BMDMs infected with Chicago4 for 4 h was assessed using ELISA. All violin plots are expressed as median $\pm$ interquartile range. Statistical analysis was performed using Kruskal-Wallis test followed by Dunn's multiple comparison test. For statistical significance, a p-value less than 0.05 was considered significant.

### Supplementary Table

**Table S1.** ***Candidozyma auris*** **strains used in this study**

| # | Organism | Strain | Other Designation | Biosample Accession | Clade | Isolation site | Reference |
|---|---|---|---|---|---|---|---|
| 1 | *C. auris* | PWT | NCCPF 470296 | | II | Pus/ Wound | [88] |
| 2 | *C. auris* | CAU-01 | AR0381 | SAMN05379608 | II | Ear canal | CDC AR Panel |
| 3 | *C. auris* | CAU-02 | AR0382 | SAMN18754596 | II | Wound | CDC AR Panel |
| 4 | *C. auris* | CAU-03 | AR0383 | SAMN05379609 | III | Blood | CDC AR Panel |
| 5 | *C. auris* | CAU-05 | AR0385 | SAMN05379620 | IV | Blood | CDC AR Panel |
| 6 | *C. auris* | CAU-07 | AR0387 | SAMN05379624 | I | Blood | CDC AR Panel |
| 7 | *C. auris* | CAU-09 | AR0389 | SAMN18754597 | I | BAL[$] | CDC AR Panel |
| 8 | *C. auris* | Chicago4 | T0318 | PRJNA1263008 | IV | | [34] |

[$] Bronchoalveolar lavage

**Table S2. Fluorescently labelled antibodies used in this study**

| Antibody/Fluorescent Probes | Clone | Vendor | Catalog No. |
|---|---|---|---|
| Propidium Iodide (Live/Dead) | | ThermoFisher Scientific | P3566 |
| CD3-BUV395 | 17A2 | ThermoFisher Scientific | 363-0032-82 |
| CD3-PE-Cy7 | 17A2 | ThermoFisher Scientific | 25-0032-82 |
| CD4-PE-Cy7 | GK1.5 | ThermoFisher Scientific | 25-0041-82 |
| CD4-Alexa Fluor 700 | GK1.5 | ThermoFisher Scientific | 56-0041-82 |
| CD11b-APC | M1/70 | ThermoFisher Scientific | 17-0112-82 |
| Ly6G-FITC | 1A8 | ThermoFisher Scientific | 11-9668-82 |
| F4/80-PE | BM8 | ThermoFisher Scientific | 12-4801-80 |
| CD11c-PE-Cy7 | N418 | BioLegend | 117318 |
| CD369-PE (Dectin-1/CLEC7A) | RH1 | BioLegend | 144304 |
| TLR4-PE-Cy7 | MTS510 | BioLegend | 117610 |
| TLR2-APC | QA16A01 | BioLegend | 153006 |
| Goat Anti Mouse IgG AF-488 | | ThermoFisher Scientific | A-24920 |
| Compensation Beads | | BioLegend | 424602 |

**Table S3. qPCR Primers used in this study**

| Species | Gene | Forward Primer | Reverse Primer |
|---|---|---|---|
| Mouse | ***Clec7a*** | CCAGCTAGGTGCTCATCTACTG | CCTTCACTCTGATTGCGGGAAAG |
| | ***Clec4n*** | CTGCTTCAGTGAAGGGACTATGG | CACTGGTGCTCCAGAAGTTCTC |
| | ***Tlr2*** | ACAGCAAGGTCTTCCTGGTTCC | GCTCCCTTACAGGCTGAGTTCT |
| | ***Tlr4*** | AGCTTCTCCAATTTTTCAGAACTTC | TGAGAGGTGGTGTAAGCCATGC |
| | ***Nox1*** | CTCCAGCCTATCTCATCCTGAG | AGTGGCAATCACTCCAGTAAGGC |
| | ***Mmp2*** | CAAGGATGGACTCCTGGCACAT | TACTCGCCATCAGCGTTCCCAT |
| | ***Mmp3*** | CTCTGGAACCTGAGACATCACC | AGGAGTCCTGAGAGATTTGCGC |
| | ***Mmp7*** | AGGTGTGGAGTGCCAGATGTTG | CCACTACGATCCGAGGTAAGTC |
| | ***Mmp8*** | GATGCTACTACCACACTCCGTG | TAAGCAGCCTGAAGACCGTTGG |
| | ***Mmp9*** | GCTGACTACGATAAGGACGGCA | TAGTGGTGCAGGCAGAGTAGGA |
| | ***Mmp10*** | TGCTGCCTATGAGGCTCACAAC | GGAGGAAAACCGAGAGTGTGGA |
| | ***Mmp12*** | CACACTTCCCAGGAATCAAGCC | TTTGGTGACACGACGGAACAGG |
| | ***Mmp13*** | GATGACCTGTCTGAGGAAGACC | GCATTTCTCGGAGCCTGTCAAC |
| | ***Mmp14*** | GGATGGACACAGAGAACTTCGTG | CGAGAGGTAGTTCTGGGTTGAG |
| | ***Mmp19*** | AGGCACTCATGGCTCCTGTCTA | TGAGCATCTCGGTCTCTTCCTC |
| | ***Mmp28*** | CTTGCTGGACACCGAGCCAAAA | CAGTTCACCAGGCGGTAGGAAA |
| | ***Timp1*** | TCTTGGTTCCCTGGCGTACTCT | GTGAGTGTCACTCTCCAGTTTGC |
| | ***Timp2*** | AGCCAAAGCAGTGAGCGAGAAG | GCCGTGTAGATAAACTCGATGTC |
| | ***Timp3*** | AGGATGCCTTCTGCAACTCCGA | GTGTAGACCAGAGTGCCAAAGG |
| | ***Elastase*** | CAGGAACTTCGTCATGTCAGCAG | AGCCATTCTCGAAGATCCGCTG |
| | ***Cathepsin G*** | AGTCCAGAAGGGCTGAGTGCTT | GCACTGTGATGAGTTGCTGGGT |
| | ***Proteinase 3*** | CTTGATCTGCAATGGCATTCTT | GGCGAAGAAATCAGGGAACT |
| | ***MPO*** | CGTGTCAAGTGGCTGTGCCTAT | AACCAGCGTACAAAGGCACGGT |
| | ***Il1b*** | AACCAACAAGT GATATTCTCC | GATCCACACTC TCCAGCTGCA |
| | ***Il6*** | TACCACTTCACAAGTCGGAGGC | CTGCAAGTGCATCATCGTTGTTC |
| | ***Tnf-*** $\alpha$ | GGTGCCTATGTCTCAGCCTCTT | GCCATAGAACTGATGAGAGGGAG |
| | ***Hprt*** | GCAGTACAGCCCCAAAATGG | AACAAAGTCTGGCCTGTATCCA |
| *C. auris* | ***Actin*** | GTATGTGCAAGGCCCGGTTTC | TGGATTGAGCCTCATCACCG |
| | ***SCF1*** | GTGAGAGTGAGGTCGGAACG | CAGCTTCTCCTTCTGGCTCC |
| | ***PIS50263.1*** | TGAGAAAGGCCGTTATGCGT | CAACGACCGACAGACAGTCA |
| | ***PIS50650.1*** | CTACAACCAAAAGCGGCACC | GTTTCTGTGGTGGATCCCGT |
| | ***XP_08167572.2*** | CGCCACAGTACGTGGCATA | TGGTGCGTCACTCATTTGT |
| | ***B9J08_004109*** | GGACAAACTGGATCTCCAGCGAA | GCAGCCAAAGAAGGCCATACTTG |
| | ***B9J08_004892*** | TGTCTCGATCGTGTCTGGTACCA | CCGCTGACACCAAGCCACATGTT |
| | ***B9J08_001531*** | GGGACATGAGCAAACACGCACCG | AACATCGCTAGGGGATGCCTGAC |
| | ***B9J08_004098*** | GCCATCCTCTTCGCTGGCTGTAT | GTCAGACTCAGTTGTAGACTGGC |
| | ***B9J08_001155*** | GTACCAGGTGGGTTCAGTCG | CCAGATGGCCTGCTCTTCAT |
| | ***B9J08_004100*** | CGTCTGAGAGCTCGTCATCGTCT | AGGATGGAGGCTCCGAAGAAGAT |
| | ***B9J08_004451*** | GGCTACCTCGCTCGCTTTAT | GCCGATGTTCAAGAGGGTGA |
| | ***B9J08_004110*** | TACGTCACCTCCGAAAACGG | GAAGGTGCCAATCGAGTCCA |
| | ***B9J08_000675*** | GCAGAAGATCAACCTGCTCCAACA | TTGATGCTCTCTTGCACTTGGCC |

208 **Table S4. *C. auris* dissemination in kidney**

| Days post inoculation | 3 | 7 | 21 |
|---|---|---|---|
| **Groups** | **Mice with dissemination / total** | | |
| Immunocompetent | 0/5 | 0/5 | 0/5 |
| Immunosuppressed | 0/5 | 0/5 | 0/5 |
| Diabetes/Ketoacidosis | 0/5 | 0/5 | 3/4 |

209

210
